# Development and Characterization of Triazole-Based WDR5 Inhibitors for the Treatment of Glioblastoma

**DOI:** 10.1101/2025.07.29.667410

**Authors:** Jesse A. Coker, Steven R. Martinez, Sang Hoon Han, Anthony R. Sloan, Amit Kumar Gupta, George Bukenya, Paul Polzer, James H. Ramos, Emma Rico, Annabella Rico, A. Abigail Lindsey, Tanvi Navadgi, Natalie Reitz, Todd Romigh, Christopher M. Goins, Chris G. Hubert, Nancy S. Wang, Feixiong Cheng, Joseph Alvarado, Samuel A. Sprowls, Justin D. Lathia, Shaun R. Stauffer

## Abstract

Glioblastoma (GBM) cancer stem cells (CSCs) contribute to tumor recurrence, treatment resistance, and dismal clinical outcomes. Genetic and pharmacological evidence suggests that the nuclear scaffolding protein WD-repeat containing protein 5 (WDR5) is a therapeutic vulnerability of the CSC population. However, previously reported WDR5 inhibitors display low permeability and are unable to penetrate the blood-brain barrier (BBB), limiting their utility in GBM. Herein, we report the structure-guided development of a novel series of triazole-based WDR5 WIN-site inhibitors designed to increase passive brain penetration. We identified triazole-based WDR5 inhibitors that are potent, passively permeable, and in some cases more brain penetrant than other scaffolds. We phenotypically assessed our novel WDR5 inhibitors in a panel of patient-derived CSC models and uncovered unique WDR5-regulated metabolic genes in GBM. We also evaluated their antiproliferative activity against CSCs both *in vitro* and *in vivo.* Finally, to identify novel combination opportunities, we screened a 2,100-compound chemical probe library and identified that the ATAD2 inhibitor **BAY-850** synergizes with WDR5 inhibitors to enhance CSC killing. Our work diversifies the chemical matter targeting WDR5, clarifies the *in vitro* consequences of WIN-site inhibition in CSCs, and encourages the future development of next-generation WDR5 inhibitors with the potential to achieve *in vivo* efficacy in the brain.

## INTRODUCTION

Despite an aggressive treatment regimen consisting of surgical resection, radiation, and chemotherapy, the prognosis for GBM, the most common primary malignant brain tumor, is dismal: median survival is 14-16 months for patients eligible for clinical trials, and 5- year overall survival is less than 3%.^1–3^ There remains an urgent need for next-generation targets and treatment paradigms that specifically address the underlying challenges of GBM. These include innate therapeutic resistance, pronounced tumor heterogeneity and cellular plasticity, and the persistence of self-renewing cancer stem cells (CSCs) that drive rapid tumor recurrence and patient mortality.^4–10^ CSCs display enhanced DNA damage response, hyper-proliferative capacity, resistance to conventional chemotherapies, and adaptive enrichment in specific hypoxic, tumorigenic niches of GBM.^11^ A growing body of evidence supports the hypothesis that specifically targeting self-renewing CSCs via novel therapeutic mechanisms could improve GBM patient outcomes.^12–15^

WDR5 is a core scaffolding component of the “WRAD” complex, which is responsible for positioning MLL-family histone methyltransferases for deposition of the activating H3K4Me^3^ epigenetic mark.^16–23^ We previously discovered that WDR5 is functionally important for CSC maintenance in GBM: genetic deletion or pharmacological inhibition of WDR5 reduced the viability and self-renewal capacity of the CSC population.^24^ These findings align with other reports demonstrating that DPY30 and RBBP5, two other WRAD complex members, are genetic dependencies in CSCs.^25, 26^ Independently of H3K4Me^3^ levels, small-molecule inhibitors of WDR5’s WDR5-interacting- (WIN-) site disrupt the protein-protein interaction (PPI) between WDR5 and MLL1, evict WDR5 from chromatin, reduce MYC-target gene expression, downregulate a core set of 50-100 ribosomal protein genes (RPGs), and reduce bulk cellular translation capacity.^27–33^ WIN-site inhibitors, such as the best-in-class imidazole-based compounds **C16** and **C10** (**Figure 1A**), achieve potent and robust single-agent antiproliferative activity *in vitro* and *in vivo* against MLL-rearranged (MLLr) leukemias that depend on oncogenic amplification of MLL1.^34–37^ Beyond these relatively rare MLLr leukemias, WDR5 inhibition has been implicated as a promising therapeutic approach in a diverse set of hematological malignancies and solid tumors, including neuroblastoma, GBM, and rhabdoid tumors.^24, 32, 37–55^ However, despite promising *in vitro* activity, there are still no reports of a potent, brain-penetrant WDR5 inhibitor, preventing *in vivo* proof-of-concept of WDR5 inhibition as a therapeutic strategy in any neurooncological indication.^56, 57^

**Figure 1.**
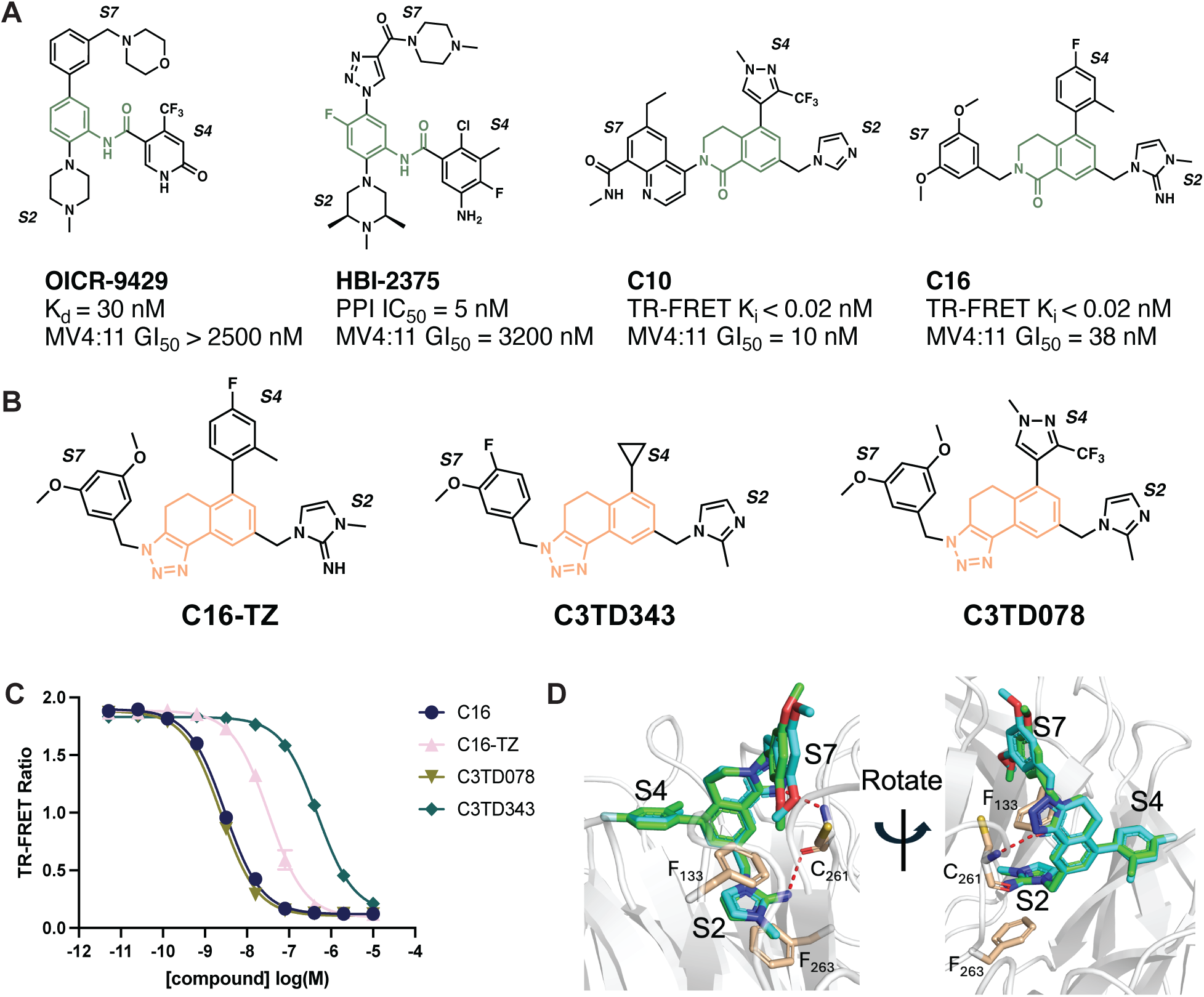
**(A)** Structures and potencies of previously reported WDR5 WIN-site inhibitors. **(B)** Structures of all novel WIN-site inhibitors profiled herein. **(C)** Representative data for the biochemical TR-FRET assay measuring compound-dependent displacement of a labelled WIN-site peptide from human WDR5. Representative data from a single replicate performed in technical quadruplicate are presented as mean +/- SD. **(D)** Overlay of WDR5:**C16** (green; PDB: 6UCS) and WDR5:**C16-TZ** (cyan, PDB: 9NCW) co-crystal structures reveals a conserved binding mode at the WIN-site, including especially a bidentate J-bond to Cys261.

Our goal was to generate potent, brain-penetrant WDR5 WIN-site inhibitors by re-engineering the core dihydroisoquinolinone scaffold of previously reported compounds like **C16.** We were inspired to remove **C16**’s lactam, reported to be a liability for BBB penetration in the ß-secretase (BACE-1) field, and replace it with a fused heterocyclic system to improve permeability. Herein, we describe a small series of WDR5 inhibitors containing a tricyclic triazole as a core amide isostere. These triazole-based inhibitors potently inhibit WDR5, maintain a conserved binding mode in the WIN-site, exhibit passive permeability, and, in some cases, improve brain penetration.

To further assess the translational potential of WIN-site inhibition in GBM, we first demonstrated that these triazole analogs potently engage WDR5 in patient-derived (PD) CSC models, and then we characterized their mechanism of action and potential for use in combinatorial regimens. Our efforts revealed that WIN-site inhibitors downregulate unique metabolic transcripts in CSCs and can be combined with the ATAD2 inhibitor **BAY- 850** to enhance CSC killing. Although significant mechanistic insights were gained *in vitro* using these tool compounds, in both intracranial and flank murine xenografts our most advanced triazoles did not reduce tumor growth or improve overall survival, likely due to insufficient potency, brain exposure, and/or metabolic stability. Our work reveals novel GBM-specific effects of WIN-site inhibition, diversifies the relatively limited chemical matter against WDR5, and provides proof-of-concept that WDR5 inhibitors can indeed be redesigned to display improved CNS penetration.

## RESULTS

### Design and Characterization of Tricyclic Triazole-Based WIN-site Inhibitors

Previously reported WDR5 WIN-site inhibitors contain a tri-substituted amide core with an Arg-mimicking “S2” group, which penetrates to the bottom of the WD40-domain central cavity, and two additional “S4” and “S7” substituents that make interactions with pockets along the surface of WDR5 (chemical structures in **Figure 1A**, binding pose in **Figure 1D**). The only brain-penetrant WDR5 inhibitor reported to date is **HBI-2375**,^56^ a methylpiperazine-S2-containing WIN-site binder disclosed in a patent owned by Huyabio (**Figure 1A**).^58^ However, **HBI-2375** is not potent, displaying a GI_50_ > 3 μM against the sensitive MLLr AML cell line MV4:11. Therefore, we initiated our medicinal chemistry campaign from a more potent starting point, namely **C16** and **C10** (**Figure 1A**), which incorporate an imidazole-based S2 discovered with fragment-based methods by Fesik and colleagues.^31, 34, 35, 37, 59^ **C16** and **C10** were reported to have a direct WDR5 K_i_ < 20 pM and an MV4:11 GI_50_ = 38 nM and 10 nM, respectively (**Figure 1A**).^37^ However, our *in vitro* profiling of C16 indicated poor passive permeability (**Table 1**), and exploratory *in vivo* rodent experiments confirmed that C16 could not penetrate the BBB (brain:plasma ratio (P:B) < 0.05; see **Supplementary Figure 1**).

To reengineer C16 into a brain-penetrant compound, we identified the core dihydroisoquinolinone as a potential moiety restricting passive permeability. Compounds like **C16** incorporate a 1,3,5-trisubstited isophthalate scaffold, which was heavily investigated in the ß-secretase (BACE-1) field for Alzheimer’s disease. Despite significant efforts, only a select few isophthalates were shown to have any measurable brain penetration.^60–62^ Based on reports that an oxadiazole bioisostere could improve the BBB permeability of such BACE-1 inhibitor scaffolds, we were inspired to replace the lactam of **C16** with a fused heterocycle.^62–64^ Our approach centered on constraining the *N*- benzylic lactam within a fused ring system, extending a flexible vector towards the S7 pocket, and maintaining a key backbone hydrogen bond between the core and Cys261. We identified a tricyclic triazole as a promising scaffold that removed the central amide whilst retaining excellent on-target potency (**Figure 1B** and **1C**).

To understand the relative properties of triazole- versus dihydroisoquinolinone-based WIN-site inhibitors, we synthesized the matched pair of **C16**, **C16-TZ** (**Figure 1B**). We characterized this pair in a biochemical TR-FRET assay, which measures the displacement of a labelled WIN-peptide from recombinant human WDR5 (**Figure 1C**).

With an imidazole-imine S2, the triazole **C16-TZ** was sub-nanomolar (K_i_ = 0.37 nM); however, this represented a 12-fold potency loss versus **C16**, which demonstrated picomolar activity (K_i_ = 0.03 nM) consistent with previously reported values.^37^ Guided by prior S2 SAR from Fesik and colleagues, we synthesized analogs with a 2-methyl imidazole S2 and a 3-methoxy phenyl S7 (**Figure 1B**) that were predicted to have improved permeability. We prepared analogs with two distinct S4 groups: a low molecular weight cyclopropyl (**C3TD343**, K_i_ = 5.9 nM) and an elaborated trifluoromethyl pyrazole (**C3TD078**, K_i_ = 0.030 nM, **Figure 1B**). As expected, the congener **C3TD078** was found to be more potent, with an equivalent K_i_ to the comparator **C16** (**Figure 1C**).

To confirm the structural basis of the triazole isostere, we solved a high-resolution 1.7 Å crystal structure of WDR5 bound to **C16-TZ** and validated a conserved binding pose matching **C16** at the WIN-site (PDB: 9NCW, **Supplemental Figure 2**). **C16** forms a bidentate hydrogen bond with the backbone of Cys261, while the imidazole-imine S2 group π-π stacks between the aromatic sidechains of Phe133 and Phe263. **Figure 1D** shows the overlay of **C16** (PDB:6UCS) with our crystal structure of **C16-TZ,** wherein the tricyclic triazole core of **C16-TZ** maintains **C16**’s bidentate hydrogen bond with Cys261 as well as the relative orientations of the S2/S4/S7 substituents. We further validated this binding pose with two other high-resolution co-crystal structures of triazole-based WIN-site inhibitors containing methyl-imidazole S2s (PDB: 9NCT, 9NCV, **Supplemental Figure 3**). Overall, **C16-TZ**, **C3TD078**, and **C3TD343** represent a diverse set of tool compounds to study the pharmacological disposition of the triazoles and characterize their potential as a novel therapeutic for GBM.

### DMPK Profiling of Triazole-Based WDR5 Inhibitors

The calculated molecular properties and DMPK profile for triazole-based inhibitors **C16- TZ**, **C3TD343**, and **C3TD078** were directly compared to the matched dihydroisoquinoline **C16** (**Table 1**). Compounds with an imidazole-imine S2, such as **C16** and **C16-TZ**, displayed low permeability in MDCK cells, high plasma protein binding (>99%), and moderate to high clearance in rat and human microsomes. The triazole analogs **C3TD343** and **C3TD078** were found to have favorable permeability (MDCK P_app_ > 2) and measurable free fraction of ∼2%. Like **C16** and **C16-TZ**, triazole-containing inhibitors consistently exhibited poor metabolic stability *in vitro* (CL_int_ > 300 mL/min/kg). Next, we moved forward with *in vivo* testing to assess the BBB permeability of triazole-based WDR5 inhibitors that displayed enhanced MDCK permeability. We first characterized the passive permeability of WDR5 inhibitors in two-minute *in situ* brain perfusion experiments, in which compounds were directly perfused into murine hearts following ligation of the descending aorta and severing of the right ventricle. After measuring the total amount of compound entering the brain at short time points, the instantaneous rate of BBB passage was quantified as a unidirectional brain uptake constant (K_in_). **C16** did not rapidly enter the brain in this perfusion setup, with a K_in_ < 5 x 10^-4^ mL/s/g (**Figure 2A**). Encouragingly, however, the triazole **C3TD343** was readily brain penetrant and displayed an order of magnitude higher rate of permeation than **C16** (K_in_ > 50 x 10^-4^ mL/s/g, **Figure 2A**).

**Figure 2.**
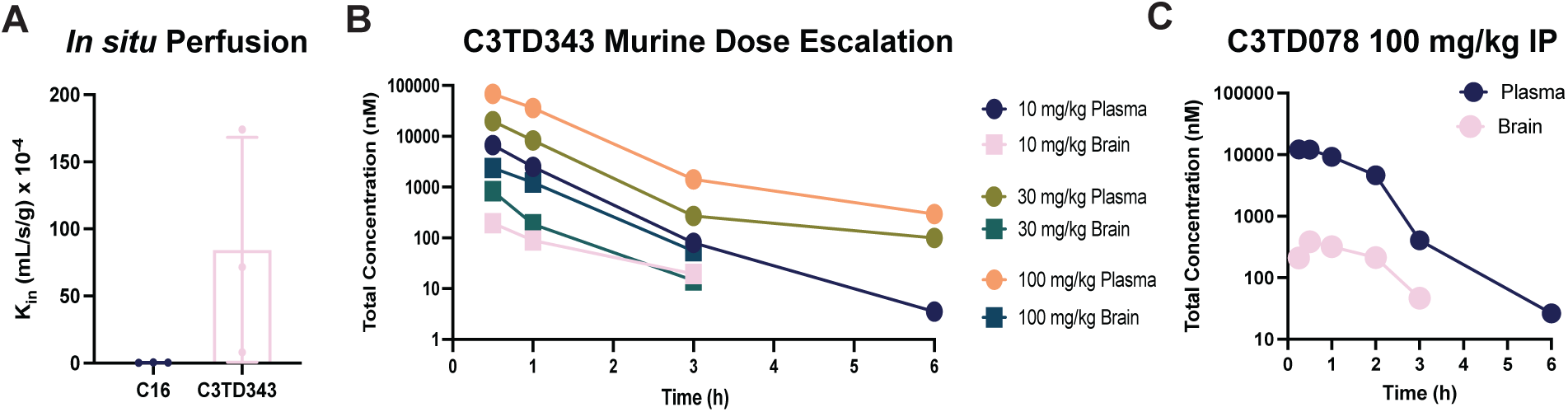
**(A)** Unidirectional brain uptake constants for **C16** and **C3TD343** in murine brain perfusion experiments (n = 3) **(B) C3TD343** total plasma (circles) and brain (squares) exposure after IP dose escalation from 10 to 100 mg/kg in mice. **(C)** Total plasma and brain exposure of **C3TD078** following a single 100 mg/kg IP dose in mice.

We proceeded with a dose-escalation experiment of **C3TD343** in mice, which revealed a dose-proportional increase in brain exposure after IP administration using 10, 30, and 100 mg/kg (**Figure 2B**). At 100 mg/kg, **C3TD343** achieved a total brain C_max_ = 2.4 μM and AUC = 2.9 μM*hrs, yielding a moderate P:B = 0.25. In agreement with the *in vitro* CL_int_ values (**Table 1**), **C3TD343** was rapidly cleared from circulation, leading to less than four hours of brain exposure. The extremely fast intake kinetics but moderate equilibrium P:B ratio for **C3TD343** was indicative of active export, and indeed we found that **C3TD343** was strongly effluxed by P-glycoprotein (Pgp *aka* MDR1) in MDR1-MDCK permeability studies (**Supplemental Figure 4**). In summary, some triazole-based WDR5 inhibitors like **C3TD343** are quite passively permeable, but their total brain exposure was limited by Pgp-mediated export and rapid hepatic clearance.

**C3TD343** contains a truncated cyclopropyl S4 substituent which confers better brain penetration but compromises potency. Compounds with an elaborated S4, such as the CF_3_ pyrazole in **C3TD078**, were up to 200-fold more potent than **C3TD343** (K_i_ = 0.03 nM vs. 5.9 nM, **Table 1**). Unfortunately, the addition of polar surface area and hydrogen-bond donors within the S4 region compromised the brain penetration of all triazoles with picomolar potency. When dosed in mice at 100 mg/kg by IP, **C3TD078** was markedly less brain penetrant (brain C_max_ = 406 nM, P:B < 0.05, **Figure 2C**) than **C3TD343**. Therefore, while we succeeded in synthesizing triazole-based WDR5 inhibitors with K_i_ < 40 pM, such potent compounds did not match the brain penetration of the moderately potent compound **C3TD343.**

### Transcriptional Profiling of WDR5 Inhibitors in GBM

Aligned with previous reports,^33, 38^ we treated L0 and DI318 PD CSCs for 72 hours with a non-cytotoxic dose (200 nM) of **C3TD078** or **C16** and then profiled the transcriptional responses by bulk RNAseq (**Supplemental Figure 5**). We observed a strong overlap between triazole- and dihydroisoquinolinone-based WDR5 inhibitors: in L0 cells, 91/102 (89%) differentially expressed genes (DEGs; log_2_FC < -0.5 & p < 0.01) from the **C16** treatment were also downregulated by **C3TD078** (**Figure 3A**); in DI318 cells, 118/150 (79%) of the **C16** DEGs were also downregulated by **C3TD078** (**Figure 3B**). Gene-set enrichment analysis (GSEA) against Howard *et al.*’s set of MV4:11 **C16**-regulated genes revealed strong downregulation of the gene set by **C16** in both L0 (NES = -2.0, FDR = 0.0, **Figure 3B**) and DI318 CSCs (NES = -2.5, FDR = 0.0, **Figure 3C**), confirming that **C16** affects a consistent set of genes across CSCs and MV4:11 leukemia cells.^33, 65^ This overlap was driven primarily by known WDR5-regulated RPGs, which were downregulated at least 2-fold by WIN-site inhibitors in GSCs (**Figure 3D**). In total, we observed n = 43 protein-coding genes that were downregulated by both **C16** and **C3TD078** in both CSC lines and can be considered the “core” set of WIN-site dependent GSC transcripts (**Supplemental Figure 6**). The most significantly enriched GO term amongst the DEGs was “cytoplasmic translation” (GO:0002181; p < 2 x 10^-16^), and Reactome pathway analysis using the “core” set of 43 genes revealed strong enrichment of “translation”, “rRNA processing”, and “peptide chain elongation” (p < 3.3 x 10^-16^, **Supplemental Figure 7**). Therefore, even in the novel cellular context of GSCs, our RNAseq data are parsimonious with the hypothesis that WIN-site inhibitors act primarily as translational stressors that deplete the ribosomal inventory.^29–32, 38^ Unlike in MLL- rearranged leukemias, we did not observe consistent downregulation of *RPL22L1* (**Figure 3D**), nor did we see consistent activation of p53 signaling (only **C16**, but not **C3TD078**, weakly enriched GO terms related to p53, see **Supplemental Figure 5**).^33^

**Figure 3.**
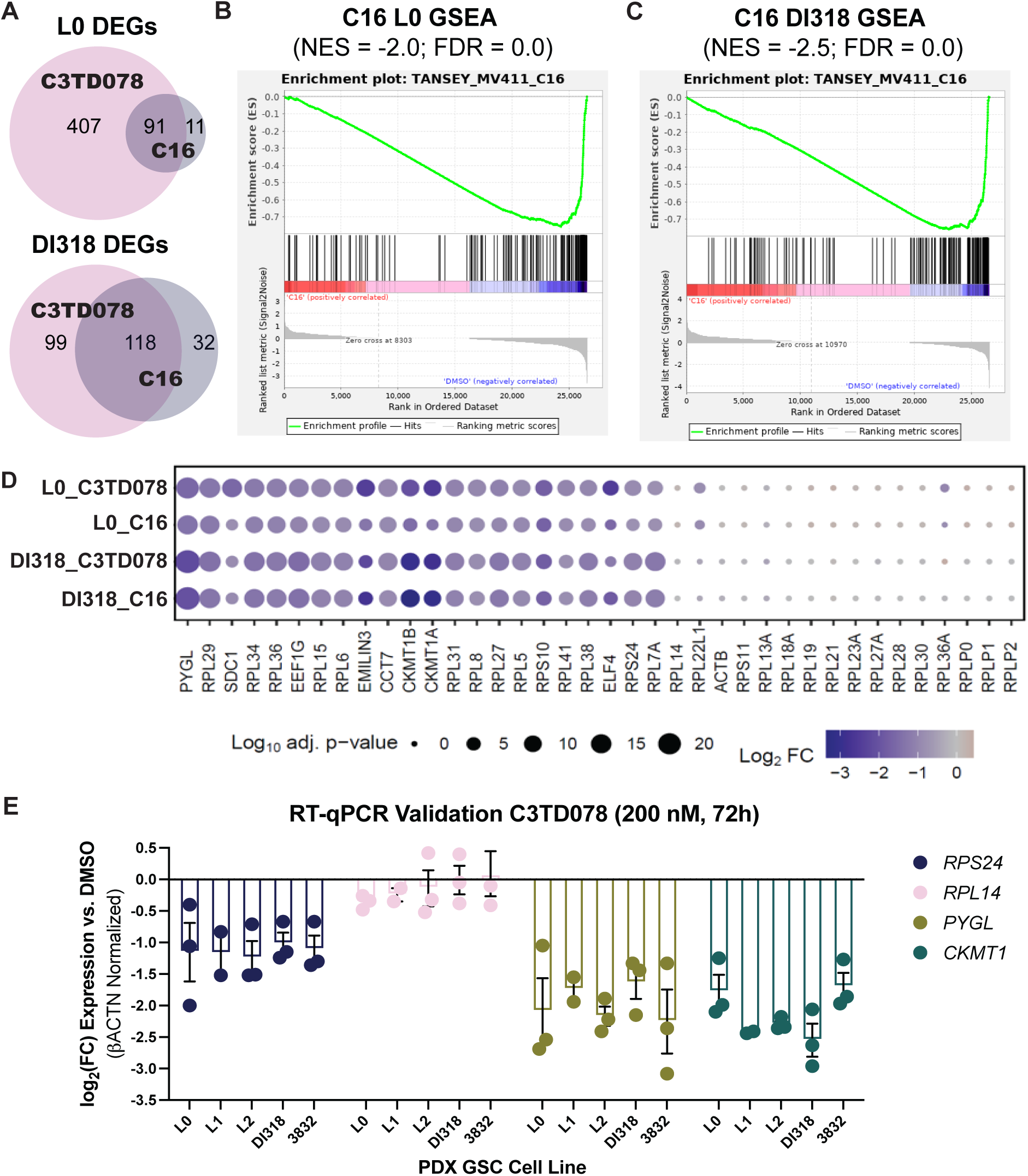
Transcriptional profiling of CSCs treated for 72 hours with 200 nM of different WIN-site inhibitors. **(A)** Overlap of differentially downregulated genes (log_2_FC < -0.5, p < 0.01) by **C16** and **C3TD078** treatment in L0 (top) and DI318 (bottom) CSCs as measured by bulk RNAseq. **(B)** GSEA against the set of genes downregulated by C16 in MV4:11 leukemia cells, as reported by Howard et al., for **C16** in L0 CSCs and **(C)** in DI318 CSCs. **(D)** Dotplot highlighting a representative set of n = 20 protein-coding genes observed to be downregulated (left) or unaffected (right) by both **C16** and **C3TD078** in both L0 and DI318 CSCs. **(E)** RT-qPCR validation for the WIN-site dependent genes *RPS24*, *PYGL*, and *CKMT1* as well as the WIN-site independent gene *RPL14*. Expression (log_2_FC) relative to DMSO is reported as mean +/- SEM for n = 2 (L1) or n = 3 (L0, L2, DI318, 3832) biological replicates.

In addition to changes in conserved WDR5-regulated RPGs, we identified n = 8 genes that were downregulated in CSCs but not in MV4:11 leukemia cells. These included two additional RPGs (*EEF1G, RPS18*), a transcription factor (*ELF4*), and two extracellular matrix proteins (*SDC1, EMILIN3*). Most intriguingly, we also observed a GBM-specific metabolic phenotype of WIN-site inhibitors, with *PYGL* (encoding the liver isoform of glycogen phosphorylase) and *CKMT1A/1B* (both encoding the mitochondrial creatine kinase CKMT1) being strongly and consistently downregulated (**Figure 3D**). CKMT1 synthesizes phosphocreatine, a metabolite that is upregulated in CSCs and that directly drives oncogenic epigenetic reprograming.^66^ *PYGL* was the single most significantly downregulated gene across both compounds in both CSC lines despite being unaffected by **C16** in MV4:11 leukemia cells (**Supplemental Figure 5**).^33^ The “liver” PYGL isoform, which performs the rate-limiting step of the cytosolic glycogen-shunt that mobilizes glucose to fuel cancer cell growth,^67, 68^ is known to be overexpressed in GBM, and genetic deletion of PYGL has been shown to enhance GBM radiosensitivity.^69^ We did observe a modest enhancement in the antiproliferative activity of **C3TD078** following 2 Gy irradiation, but only in a single CSC line (DI318; **Supplemental Figure 8**).

We extended these RNAseq observations by RT-qPCR validation in five different PD CSC models (**Figure 3E**). The WDR5-bound, WIN-site responsive RPG, *RPS24*, was downregulated ∼2-fold by **C3TD078** treatment in all five models, while the WDR5- independent RPG *RPL14* was unaffected, as expected and reported by others.^31^ In agreement with our RNAseq data, both *PYGL* and *CKMT1* were downregulated by ∼4-fold in all five CSC models (**Figure 3E**). We also validated that the “liver” *PYGL* isoform is expressed at >1,000 fold higher levels than the “brain” *PYGB* isoform in CSCs, and we found that WIN-site inhibitors had no effect on the already low expression of *PYGB* (**Supplemental Figure 9**). Taken together, our transcriptional profiling revealed both ubiquitous RPGs and context-specific metabolic transcripts (*PYGL*/*CKMT1*) that were downregulated by WIN-site inhibition in CSCs.

### Target Engagement of WDR5 Inhibitors in CSCs

To confirm intracellular target engagement of our WDR5 inhibitors in CSCs, we first used a cellular thermal shift (CETSA) assay in intact L0 CSCs. 10 μM of **C16** induced a τιT_m_ = +35 °C in L0 cells but did not change the T_m_ of βACTN (**Figure 4A**). For intracellular K_d_ determinations, we performed compound titrations (two-hour pre-treatment) with an isothermal melt at 70 °C followed by centrifugation to separate ligand-bound from apo WDR5. Representative data for a potent compound (**C3TD078**, K_d_ = 2 nM) and a weak compound (**C3TD343**, K_d_ = 200 nM) are provided in **Figure 4B**. Across all tested triazoles (n = 17), we observed a linear correlation (R^2^ = 0.89) between the biochemical K_i_ as determined by TR-FRET and the intracellular K_d_ as determined by CETSA (**Figure 4C**). We also deployed the CETSA in washout experiments to establish the profoundly long off-rate of **C3TD078**, which remained bound to WDR5 for at least 21 hours following compound removal (**Figure 4D**). C16 also remained bound to WDR5 for > 20 hours following washout (**Supplemental Figure 10**), suggesting that, across chemotypes, potent WDR5 inhibitors display a slow intracellular off-rate.

**Figure 4.**
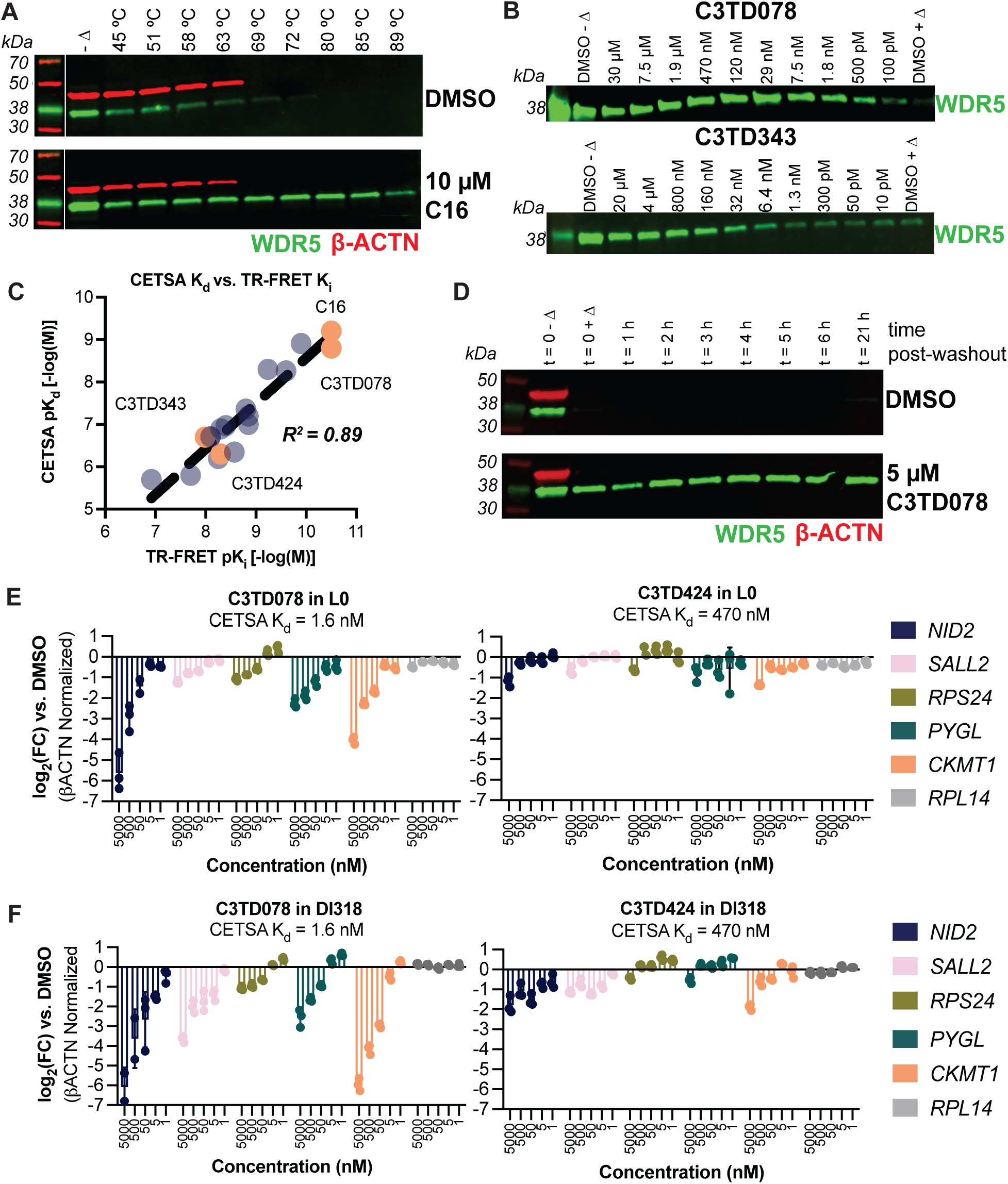
**(A)** Melting curve for WDR5 (green) and ACTB (red) determined by CETSA-WB following a two hour treatment of L0 CSCs with either DMSO (top) or 10 μM **C16** (bottom). **(B)** Isothermal CETSA-WB (70°C) for L0 CSCs treated for two hours with varying doses of the potent binder **C3TD078** (top) or the weak binder **C3TD343** (bottom). **(C)** Correlation between biochemical potency (TR-FRET K_i_) and cellular potency (L0 CETSA K_d_) as determined for n = 17 different WDR5 inhibitors. **(D)** Washout experiment following the treatment of L0 CSCs for two hours with either DMSO (top) or 5 μM **C3TD078** (bottom) followed by compound washout. Samples were prepared for isothermal CETSA-WB at the indicated timepoints post-washout. **(E)** RT-qPCR analysis for the indicated genes after treating L0 CSCs with the indicated doses of the potent inhibitor **C3TD078** (left) or the weak inhibitor **C3TD424** (right) for 72 hours. Data are presented as mean +/- SD from a single biological replicate completed in technical triplicate. **(F)** Same as in (E) but with DI318 CSCs.

To further validate the on-target nature of the transcriptional effects observed in **Figure 3**, we performed head-to-head dose-response experiments by RT-qPCR using both a potent (**C3TD078,** CETSA K_d_ = 1.6 nM) and weak WDR5 inhibitor (**C3TD424**, CETSA K_d_ = 500 nM, see **Supplemental Figure 11** for chemical structure). In addition to the RPGs and metabolic transcripts identified by RNAseq in **Figure 3**, we also quantified the expression of *SALL2* and *NID2*, which were identified as WDR5-bound genes in CSCs by CHIPseq in our previous study.^24^ In both L0 and DI318 cells, **C3TD078** (CETSA K_d_ = 1.6 nM) achieved potent, dose-dependent inhibition of *RPS24*, *PYGL*, *CKMT1*, *SALL2,* and *NID2*, while the less potent **C3TD424** showed only modest inhibition of these genes at only the highest doses (**Figure 4E/4F**). As expected, neither compound impacted the expression of the WDR5-independent gene *RPL14*. Across both cell lines and all five genes, the average IC_50_ = 80 nM for **C3TD078**-induced transcriptional downregulation, which closely matches reported IC_50_ values for **C16**’s transcriptional activity in MLLr leukemia cells.^38^ Taken together, these RT-qPCR data align with the CETSA and support potent on-target inhibition of WDR5 by triazole-based WIN-site inhibitors in CSCs.

### Anti-Proliferative Activity of Triazole-Based WDR5 Inhibitors Against CSCs

In MLLr-leukemia cell lines like MV4:11, potent WIN-site inhibitors like **C16** achieve low double-digit nM antiproliferative activity, while in insensitive cell lines, like A549, the same WIN-site inhibitors display GI_50_ > 2 μM.^35, 59^ In a 7-day assay using five different CSC models, neither **C16** nor **C3TD078** achieved remarkable antiproliferative activity, with GI_50_ values between 1—5 μM (**Figure 5A**). There was a trend towards more potent GI_50_‘s with **C3TD078**/**C16** relative to the weaker compound **C3TD343**, suggesting a modicum of on-target, WDR5-mediated antiproliferative activity across this CSC panel. However, we also observed that both **C16** and **C3TD078**, likely due to their protonatable S2 warheads, induced phospholipidosis (PLD) and disrupted cellular membrane architecture at doses above 1 μM (**Supplemental Figure 12**). Therefore, we could not rule out the possibility of off-target PLD dominating the observed antiproliferative phenotype.^70–72^

**Figure 5.**
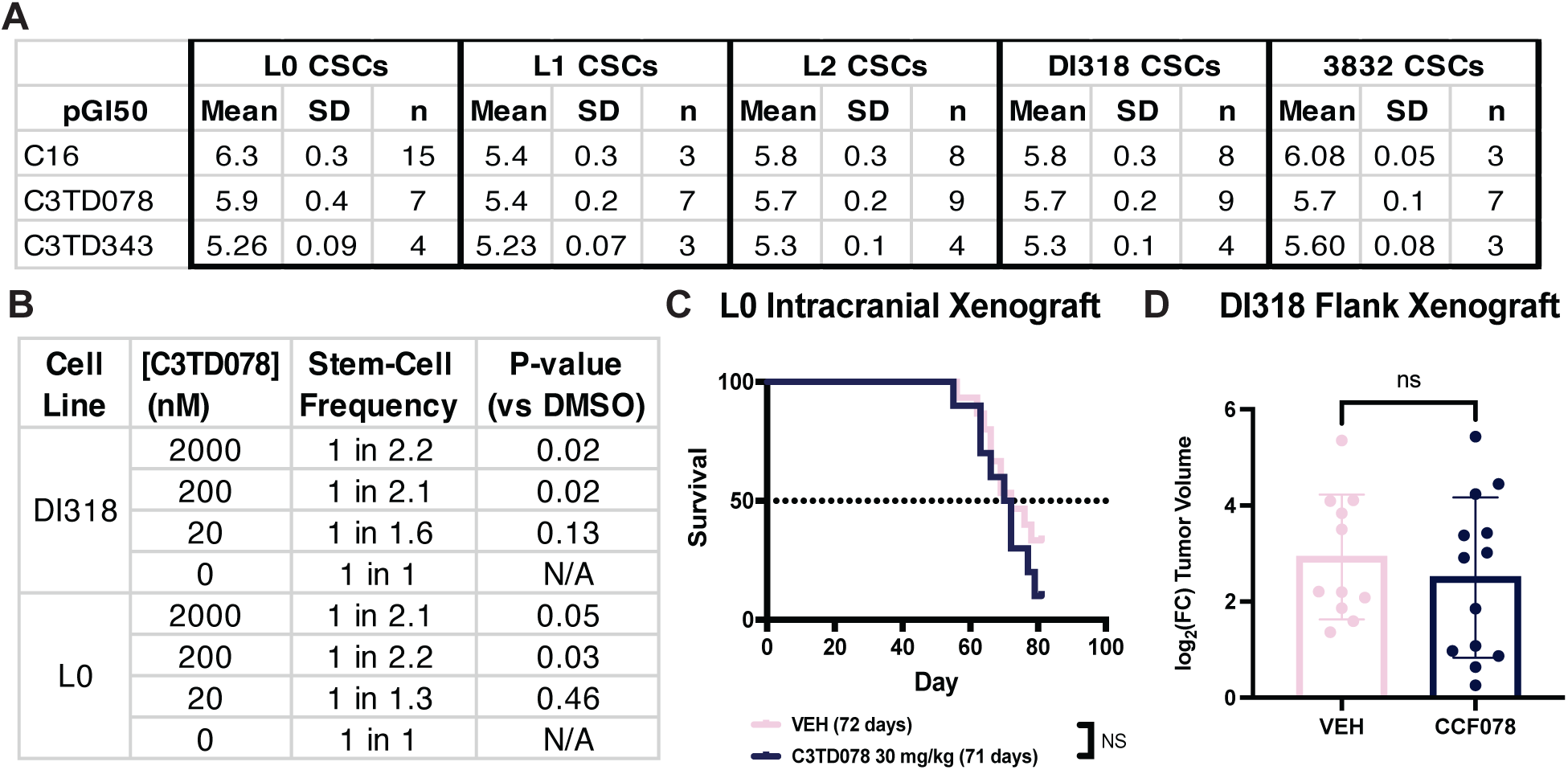
**(A)** Anti-proliferative activity of WIN-site inhibitors against n = 5 CSC lines in a 7-day CTG® viability assay. **(B)** Quantification of ELDA after treatment of DI318 or L0 CSCs for 14 days with the indicated doses of **C3TD078**. Stem-cell frequency and p-value relative to DMSO were calculated as previously reported.^95^ **(C)** Kaplan-Meier survival curve for mice implanted with intracranial L0 PDX tumors and treated daily with vehicle or 30 mg/kg QD IP **C3TD343**. **(D)** 7-day fold-change in tumor volume of flank DI318 PDX tumors following treatment with vehicle or 30 mg/kg QD IP **C3TD078**. Data were combined from two separate experiments and end-point tumor-volume increase was compared by a t-test.

Next, we examined the ability of **C3TD078** to reduce the stem-cell frequency of CSCs with a traditional extreme limiting dilution assay (ELDA). We observed that **C3TD078** dose-dependently and significantly decreased the stem-cell frequency of DI318 and L0 CSCs, but only modestly—200 nM of **C3TD078** resulted in a halving of the number of stem cells (**Figure 5B**). These effects on self-renewal were much smaller than previously reported with 5 μM of **C16**, perhaps because of WDR5-independent cytotoxicity at this high dose.^24^

Finally, we executed two *in vivo* studies to assess the activity of our triazole-based WDR5 inhibitors in murine xenografts (PDX). First, we characterized our most brain penetrant but modestly potent compound **C3TD343** in an intracranial xenograft. We treated mice implanted with intracranial L0 PDX tumors for up to 80 days with 30 mg/kg QD IP **C3TD343**, which was well tolerated. However, we observed no impact on survival (**Figure 5B**). We next performed a shorter-term flank DI318 PDX model with our most potent, slow off-rate, and peripherally bioavailable triazole **C3TD078**. Even in this simplified flank model with a potent WDR5 inhibitor, **C3TD078** did not reduce tumor growth (**Figure 5C**). Taken together, our *in vitro* and *in vivo* data suggest that, despite potent cellular target engagement, WDR5 WIN-site inhibitors display unremarkable single-agent antiproliferative activity against CSCs.

### Synergy Between WDR5 and ATAD2 Inhibitors Against CSCs

Since WIN-site inhibitors did not robustly kill CSCs, we decided to pursue a combination approach, which is precedented for this mechanism of action: the BCL-2 inhibitor Venetoclax synergized with WDR5 inhibitors to kill AML cells,^36^ while the p53 activator Nutlin-3A enhanced the efficacy of WDR5 inhibitors against rhabdoid tumors.^38^ However, to our knowledge, there have been no reports of an unbiased synergy screen to identify combination opportunities from a large library. We were further encouraged to consider broad screening since we observed no synergistic efficacy between Venetoclax, the p53 activator RG7112, or other rational combinations with WDR5 inhibitors in our CSC models (**Supplemental Figure 13**). Therefore, we screened the Cleveland Clinic’s 2,100 compound Pinpoint Chemical Probe Library (PiCL),^73^ which contains FDA-approved drugs and publicly available chemical probes, for compounds that killed CSCs more effectively in the presence of the WDR5 inhibitor **C16**. We first screened the entire library (at 1 μM) in the presence of a non-cytotoxic dose (200 nM) of **C16** using a 7-day CTG® viability assay in both L0 and DI318 cells. Any treatment achieving a >80% reduction in cell viability in either cell line was then counter-screened in the presence and absence of **C16** to identify synergistic combinations. We identified 7 PiCL compounds that killed L0 cells better in the presence of 200 nM **C16** (**Figure 6A**); in DI318 cells, we identified 3 combination opportunities (**Figure 6B**). There was a single hit shared between both cell lines: **BAY-850**,^74^ an inhibitor of the bromodomain-containing, epigenetic regulator ATAD2 (ATPase family AAA domain-containing protein 2).^75^

**Figure 6.**
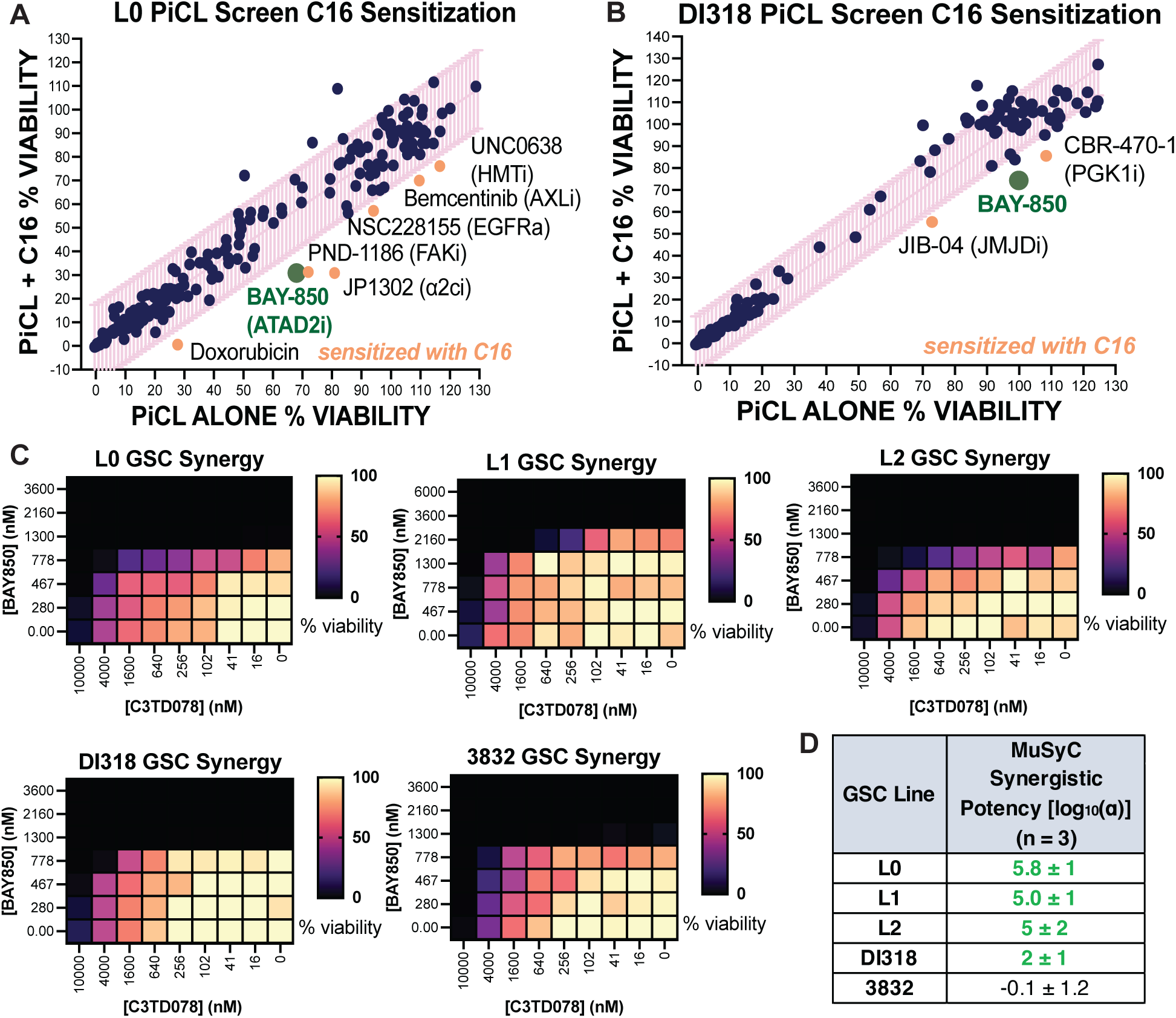
**(A)** Synergy screen with the Pinpoint Chemical Probe Library (PiCL; 1 μM) by CTG® viability after 7-day treatment in the presence (y-axis) or absence (x-axis) of 200 nM **C16**. The 95% confidence interval of the linear regression is displayed in pink. PiCL library members that reduced the viability of CSCs more in the presence of **C16** are towards the southeast of the plot and are labeled. **(B)** Same as (A), except in DI318 CSCs. **(C)** 7-day CTG® viability cross-titrations with **C3TD078** and the ATAD2 inhibitor **BAY-850** in five different CSC lines. Data are a representative of three biological replicates. **(D)** Overall quantification of the synergistic potency, as assessed by the MuSyC algorithm, between **BAY-850** and **C3TD078** inhibitors in CSCs.

After validating the expression of ATAD2 in our CSC models by western blot (**Supplemental Figure 14**), we proceeded to validate the synergy suggested by the PiCL screen in five different CSC models by performing cross titrations with **BAY-850** and **C3TD078** (**Figure 6C**). We analyzed the dose-response matrices with the MuSyC algorithm, which can disentangle synergistic efficacy (i.e. combinatorial increases in the maximum cytotoxic effect) from synergistic potency (i.e. a left-shift in the combinatorial IC_50_, denoted α).^76^ We observed a relatively steep dose-response curve with **BAY-850** alone, perhaps due to its mechanism of action as an ATAD2 dimer inducer.^74^ Nevertheless, we did observe clear synergistic potency increases, with **BAY-850** left-shifting the GI_50_ of **C3TD078** (α > 1) in 4/5 CSC models (**Figure 6D**). These data resonate with a recent report demonstrating that ATAD2 is a downstream effector of WDR5 that plays a role in the growth of ALL cells.^77^ We did not observe synergy with the ATAD2 bromodomain competitive inhibitor **GSK8814**,^78^ suggesting that the binding of ATAD2 at H4Kac is dispensable for the observed synergy (**Supplemental Figure 15**). While limited to only *in vitro* observations that require future *in vivo* validation, these data implicate ATAD2 dimerization as a previously unprecedented mechanism for combination with WDR5 WIN-site inhibitors.

## DISCUSSION

Herein, we disclose a novel series of triazole-based WDR5 inhibitors designed to be passively brain penetrant for the treatment of GBM. Mechanistic studies with our triazoles confirmed potent intracellular WDR5 target engagement, and the first reported bulk RNAseq experiment in WDR5-inhibitor-treated CSCs revealed both ubiquitous RPGs and a set of CSC-specific transcripts (*PYGL*, *CKMT1*, *SALL2*) that were downregulated. These functional data improve upon our previous work, which reported the *in vitro* activity of **C16** in CSCs at doses ≥ 3 µM,^24^ through the consistent use of lower doses (200 nM) of **C16**/**C3TD078** that do not induce off-target PLD. On exemplar triazole, **C3TD343**, showed robust passive BBB permeability, but it was not efficacious in an intracranial xenograft due to a combination of low potency, rapid hepatic clearance, and Pgp-mediated export. We also characterized the potent but only peripherally bioavailable triazole **C3TD078** in proliferation assays and a flank xenograft, but we consistently observed unremarkable CSC killing.

Our work emphasizes a considerable challenge for the field: WDR5 inhibitors must have exquisitely potent direct affinity to have any functional effects in cells.^56^ For example, **C3TD078^’^**s potency right-shifts from biochemical TR-FRET K_i_ = 30 pM, to cellular CETSA K_d_ = 1.6 nM, to functional RT-qPCR IC_50_ = 80 nM. This represents, in total, a 2,700-fold right-shift in **C3TD078**’s potency from direct WDR5 affinity to transcriptional downregulation. Therefore, maintaining both pM direct affinity (with molecules consistently displaying MW > 500 Da) and brain penetration is a difficult hurdle, especially with the requirement for a weakly basic S2 Arg-mimetic. The S4-truncated **C3TD343** does indeed cross the BBB, but like the other reportedly brain penetrant analog **HBI-2375**, it is too weak (K_i_ = 5 nM) to have pronounced functional effects in cells: a similar 2,700-fold shift would imply an unfeasible transcriptional IC_50_ > 13 µM, well above the brain exposures achievable with **C3TD343** (C_max_ = 2.4 µM at 100 mg/kg). Ultimately, we never identified a triazole with both picomolar WDR5 affinity (like **C3TD078**) and high brain exposure (like **C3TD343**).

It is difficult to establish with certainty whether the failure of **C3TD343** in our intracranial xenograft was due to the low potency of this specific compound or a broader lack of efficacy of WIN-site inhibition in GBM. The low efficacy of **C3TD078** in flank models, as well as the modest GI_50_’s in 7-day *in vitro* proliferation assays with **C16** and **C3TD078**, support the hypothesis that single-agent WDR5 inhibition is not a transformative paradigm in GBM. An opportunity to potentiate the activity of WDR5 inhibitors is cotreatment with the ATAD2 inhibitor **BAY-850**, which we identified as a synergistic combination from a 2,100 compound chemogenomic screen. However, this combination hypothesis requires validation *in vivo*.

It is well established that the biology of GBM is distinctive in the brain, likely due to specific microenvironmental cues that can only be recapitulated in the correct target tissue. It has been elegantly demonstrated that genetic knockout of DPY30, an essential member of the WRAD complex alongside WDR5, is dispensable for GBM growth *in vitro* but absolutely essential for *in vivo* intracranial GBM growth and self-renewal.^25^ Many other epigenetic proteins, including DOT1L, SETD2, BRD4, and JMJD6 only score as essential genes in genetic dropout screens conducted intracranially, while they are wholly dispensable for CSC growth in tissue culture.^79^ Therefore, because of the complex local biology and epigenetics of GBM in the brain, the therapeutic efficacy of WDR5 inhibitors can only be properly ascertained in the correct tissue microenvironment. We hypothesize, then, that the failure of **C3TD343** to enhance survival in an intracranial model may be due to low potency; it remains an open question if picomolar WIN-site inhibitors are antiproliferative in intracranial GBM models.

We are hopeful that our report will catalyze future medicinal chemistry efforts to enable the testing of WDR5 inhibitors in a biologically relevant setting. Our experience suggests this will be challenging, and will likely require the discovery of more compact, lower molecular weight WDR5 inhibitors distinct from the reported chemical space. We also recognize the potential for unique formulations and routes of administration (such as lipid nanoparticle encapsulation or transient ultrasound-mediated BBB opening) that could overcome the need for a passively penetrant compound. Finally, given that WDR5 is a genetic dependency specifically in the stem-like outer rim of GBM organoids and tumors,^24^ it merits further evaluation of the ability of WIN-site inhibitors to reduce GBM recurrence following surgical resection. Overall, our work diversifies the known chemical matter against WDR5, clarifies the *in vitro* phenotype of WIN-site inhibition in CSCs, and sets the foundation for future development of brain-active WDR5 inhibitors that display the requisite combination of picomolar potency and brain bioavailability.

## METHODS

### Sex as A Biological Variable

All *in vivo* efficacy studies were conducted with an equal number of male and female mice to capture known sex-differences in GBM survival. All *in vivo* PK experiments were conducted with only male mice, which is industry standard.

### Statistics

Relevant statistical tests and number of biological replicates are described in the corresponding figure legends.

### Study Approval

All animal procedures were performed in accordance with the guidelines and protocols approved by the Institutional Animal Care and Use Committee at the Cleveland Clinic and by the Walter and Eliza Hall Institute Animal Ethics Committee. NSG (NOD.Cg- PrkdcscidIl2rgtm1Wjl/SzJ) mice were obtained from the Biological Research Unit (BRU) at Cleveland Clinic Research.

### Primary Cell Culture

DI318 was derived from de-identified samples collected from the Cleveland Clinic Brain Tumor and Neuro-Oncology Center. 3832 were a gift from Duke University, while L0, L1, and L2 CSCs were a gift from the University of Florida. All CSC models were maintained in Neurobasal^TM^ Complete Medium [NBMc: Neurobasal Medium (Gibco #12349015) + 1X GlutaMax + 1X Sodium Pyruvate + 1X Pen/Strep + 1X Vitamin B27-A Supplement (Life Technology #12587010) + 10 μg/L EGF (R&D #236-EG) + 10 μg/L FGF2 (R&D #4114-TC)] as described previously.^24, 80^

### Purification of WDR5

Recombinant human WDR5 was cloned, expressed, and purified as previously described. ^37^ Briefly, human WDR5 residues Ser22—Cys334 were codon optimized for *E.coil* and cloned into a pET27 plasmid downstream of a SUMO-protease cleavable NT-His6-SUMO fusion. T7 Express *lys*Y/I^q^ *E.coli* cells (New England Biolabs) were transformed with the expression plasmid were grown in Terrific Broth media supplemented with Kanamycin at 37°C with shaking to an optical density λ_600nm_ = 0.6, then reduced to 30°C and induced with 1 mM isopropyl-beta-D-thiogalactoside overnight. Induced cultures were pelleted, resuspended in 1X PBS pH 7.4, 300 mM NaCl, 20 mM imidazole, 5 mM β-Me, and 10% glycerol and lysed via sonication. Lysates were clarified by centrifugation, loaded onto an equilibrated NiNTA column, washed with resuspension buffer, and eluted over a gradient of increasing [imidazole]. His-SUMO-WDR5 used in the TR-FRET assay was further purified via size exclusion chromatography using 1x PBS pH 7.4, 300 mM NaCl, 5 mM β-Me running buffer, concentrated to 1-2 mg/ml, aliquoted, and froze at -80°C for storage. Tag free WDR5 used for crystallography was dialyzed overnight with SUMO protease and passed over an NiNTA column to remove the His6-SUMO tag. Tag free WDR5 was further purified via size exclusion chromatography using 20 mM HEPES pH 7.0, 250 mM NaCl, and 5 mM DTT, concentrated to 10 mg/ml, aliquoted, and frozen at -80°C for storage.

### WDR5 Co-Crystallization and Data Processing

WDR5 at 10 mg/ml was titrated with 2-4% v/v inhibitor (10 mM DMSO stock), incubated at room temperature for 1 hour, and co-crystallized by hanging drop vapor diffusion. Crystals formed in previously reported conditions of 0.2 M ammonium acetate, 20-30% PEG 3350, and 0.1 M Bis-TRIS, TRIS, or HEPES pH 6-8 within 1-3 days of incubation at 16°C.^37^ Crystals were cryoprotected with 20% glycerol, looped, and plunge froze in liquid nitrogen. X-ray diffraction data was collected at the Advanced Photon Source LS-CAT 21- ID-G and 21-ID-F beamlines under cryogenic conditions. Diffraction data was indexed, scaled, and merged using HKL2000.^81^ Structures were phased using molecular replacement (Phaser-MR) with PDB: 6UOZ and refined using Phenix Refine with manual rounds of refinement in Coot.^82–85^ Ligand restraints were generated using ELBOW and final figures generated using PyMOL.^86^ Final structure statistics can be found in **Supplemental Figure 2**. Structure maps and coordinates were deposited within the Protein Data Bank (PDB: 9NCW, 9NCT, 9NCV).

### WDR5 TR-FRET Binding Assay

Inhibitor *K*_i_ values were measured using a previously published competitive TR-FRET assay.^37^ Compounds dissolved at 10 mM in DMSO were serial diluted 5-fold over 10 concentration points and acoustically transferred into 384-well OptiPlates (Revvity) using a Labcyte Echo 550. A 20 μL solution of 1 nM LanthaScreen^TM^ Elite Tb-anti-His antibody (ThermoFisher), 150 nM “10-mer-Thr-FAM” peptide probe (ARTEVHLRKS-(Ahx-Ahx)(Lys-(5-FAM))) synthesized by GenScript, and 2 nM recombinant His6-SUMO-WDR5 all diluted in assay buffer (1x PBS pH 7.2, 300 mM NaCl, 0.5 mM TCEP, 0.1% CHAPS) was titrated into each well of the compound-stamped plate for a final 0.1% v/v DMSO. Assay plates were centrifuged, covered, and incubated at room temperature for 1 hour prior to spectroscopic measurement (λ_ex_ = 340 nM, λ_em1_ = 495 nm, λ_em2_ = 520 nm) on a BioTek Cytation5 plate reader. The 520/495 TR-FRET ratio was calculated and plotted against the logarithmic inhibitor concentration. For *K*_i_ determination, the sigmoidal displacement curves were fit with a “One site – Fit Ki” analysis using PRISM10 and constraining “HotNM” = 150 and the “HotKdNM” = 2. The reported *K*_i_ values were calculated from at least three biological replicates completed in technical quadruplicate.

### WDR5 Live Cell CETSA

500,000 L0 CSCs, grown in suspension as spheroids, were split with Accutase® and seeded in 1 mL of NBMc into each well of a 24-well dish. Compounds were prepared in DMSO at 333X concentration and 3 μL (0.3% v/v DMSO) was added to the corresponding wells and incubated for 2 hours. Cells were harvested by centrifugation (5 min, 500 x *g*) into SafeLock tubes (Eppendorf), washed once with 400 μL of PBS, and resuspended in 25 μL of CETSA Buffer (PBS + 1X Roche cOmplete^TM^ Protease Inhibitor Cocktail + 3X Benzonase). For generation of thermal melting curves, samples were heated for three minutes at 45-89 °C. For isothermal K_d_ determination, samples were heated for three minutes at 70 °C. One DMSO-treated sample was left unheated (representing 100% abundance *i.e.* fully bound) and one DMSO-treated sample was heated to 70 °C (representing 0% abundance *i.e.* fully unbound). After heating, samples were lysed by 6X freeze-thaw cycles in LN_2_/hot water, rested for 10 min at RT, and then centrifuged for 30 minutes at >14,000 x *g* at 4 °C. 20 μL of the supernatant was transferred to a fresh tube into 5 μL of SDS-PAGE Loading Dye, boiled, and separated on 4-20% Mini-PROTEAN® Precast TGX Gels (Biorad). Gels were transferred to PVDF with a Trans-Blot Turbo (Biorad), probed with primary antibodies to WDR5 (1:1000, rb CST #13105) and ACTB (1:3,000 ms #CST3700), and amplified with 1:5,000 LI-COR goat secondary antibodies (IRDYE680RD Anti-Mouse & IRDYE800CW Anti-Rabbit). WDR5 band intensity was quantified on a LI-COR Odyssey® imager and normalized from 0% (DMSO heated) to 100% (DMSO unheated), plotted against log[compound], and midpoint K_d_ values were extracted by non-linear regression in GraphPad PRISM. For CETSA washout experiments, a 10 cm dish with 6-8 x 10^6^ L0 cells was treated for 2 hours with 5 μM compound or DMSO, washed 3X times with 15 mL of PBS (3 min, 400 x *g*), and split into 10 x 1 mL of fresh NBMc in a 24-well dish. Single wells were taken and prepared for CETSA-WB as described above at different time-points over the course of 21 hours.

### RT-qPCR and Primer Sets

PD CSC lines L0, L1, L2, DI318, and 3832, all grown as spheroids in NBMc, were split with Accutase® and 300,000 cells were seeded in 4 mL of NBMc into 6-well dishes (three wells per condition). 4 μL of a 1,000X compound stock prepared in DMSO (0.1% v/v DMSO) was added and cells were grown at 37 °C/5% CO_2_ as standard for 72 hours. Cells were harvested by centrifugation (5 min, 500 x *g*) into 2 mL Eppendorf Tubes and total RNA was purified on spin columns using the RNeasy Plus Kit (Qiagen, includes gDNA removal columns). 2.5 μg of total RNA was treated with ezDNAse^TM^ (Thermo Fisher) and then reverse transcribed with the SuperScript^TM^ IV VILO^TM^ RT Master Mix (Invitrogen). cDNA was diluted 1:100 in nuclease-free water and then transcripts were quantified using the PowerUp SYBR® Green Master Mix (Thermo Fisher) on a QuantStudio^TM^ 3 RT-qPCR machine according to the manufacturer’s recommendations. Please see the table below for primer sets. Raw Cq values were normalized per sample to the housekeeper ACTB and then the fold-change relative to DMSO treated cells was calculated as FC = 2^-ΔΔCq^. **TRANSCRIPT FORWARD PRIMER (5’-3’) REVERSE PRIMER (5’-3’)**

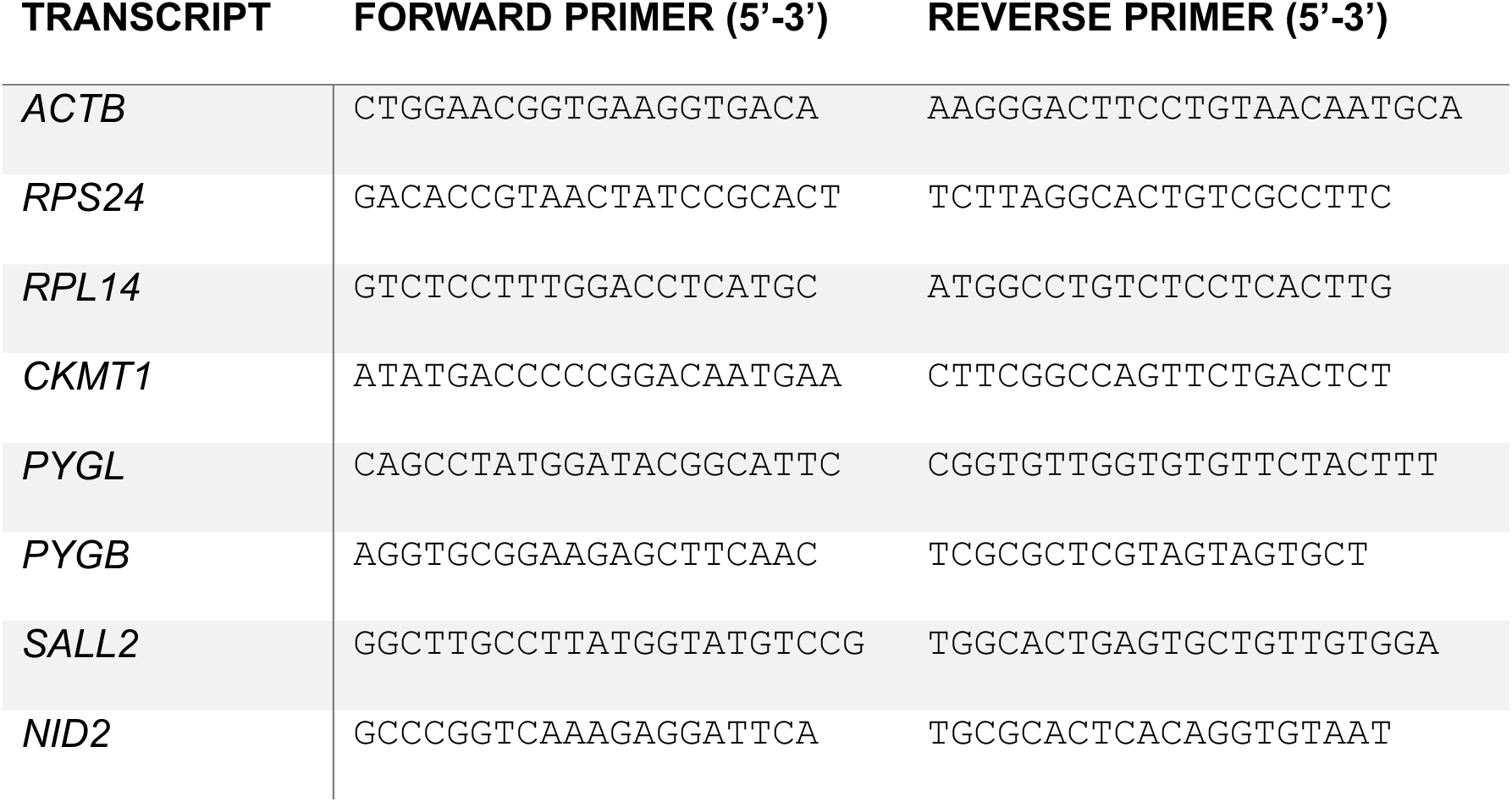

### 384w CSC Viability Assays

333X-1000X compound dilution series were prepared in 100% DMSO by serial-dilution and then stamped by acoustic dispensing (50-150 nL, 0.1-0.3% DMSO v/v) into 384-well sterile TC-treated white opaque plates using a Labcyte Echo 550. For GI50 determination, compounds were stamped in technical triplicate, while for synergy cross-titrations dose-response matrices were prepared in technical duplicate. Different CSC lines, grown in suspension as spheroids, were split with Accutase® and seeded directly on top of the pre-stamped compounds (50 μL of NBMc and 700 cells/well). The outer four Rows (A,B & O,P) and four columns (1,2 & 23,24) were filled with 80 μL of PBS to minimize evaporation. Cells were incubated with compounds for 7 days at 37 °C/5% CO_2_ prior to the addition of 25 μL/well Cell-Titer Glo® (Promega) and luminescence quantification on a Revvity EnVision^TM^ plate reader. Percent viability values were calculated by normalization to DMSO treated cells (100%) or staurosporine treated cells (0%) and GI50 values were extracted by non-linear regression in GraphPad PRISM. For the cross- titrations of two compounds, synergy scores were calculated using the MuSyC webserver.^76^ For the PiCL screen, the entire library of ∼2,100 FDA-approved drugs and chemical probes was screened by CTG® as described (except for 5 days) at 1 μM as singletons in the presence of 200 nM C16 against L0 and DI318 CSCs. Any PiCL compound achieving > 80% reduction in cell viability against either CSC line in this primary screen (n = 245) was then counter-screened in the presence or absence of 200 nM C16 to identify potential synergistic combinations.

### RNAseq and Data Analysis

L0 and DI318 CSCs were treated were treated for 72 hours with DMSO (0.1% v/v), 200 nM C16, or 200 nM C3TD078 (triplicate for all treatments) and total RNA was purified as described in the RT-qPCR section and then quality controlled for integrity and concentration on a Qubit® (Invitrogen). Bulk RNAseq was performed at MedGenome Inc. using the Illumina® Stranded mRNA Library Prep Kit and sequencing at 20M paired-end read-depth on a NovaSeq 6000 (PE100). The raw bulk RNA-seq paired end (R1-R2, 2*100 bp) sequencing fastq data was first subjected to the quality check analysis using FastQC (v0.11.9). Then, raw reads were quality filtered utilizing multithreaded fastp tool (v0.23.2) (30423086), majorly implementing deduplication, auto-detect adapter removal, N base removal, Phred quality filtering (>=Q20 is qualified), and unqualified bases percent limit (=30) with other default parameters. Further, quality filtered reads were subjected to the alignment employing Rsubread (v2.16.1) align function with human genome reference build GRCh38.^87, 88^ The mapped binary alignment map (BAM) files were indexed using samtools (v1.16.1),^89^ and gene-wise read counts were summarized applying featureCounts function from Rsubread, where inbuilt hg38 annotation was used and duplicated reads were ignored. Further, gene symbols were obtained using org.Hs.eg.db (v3.15.0) annotation package operating on ENTREZIDs from count table. Respectively, edgeR (v3.38.4) and limma-voom (v3.52.4) packages were employed to accomplish differential transcriptional regulation analysis.^90–92^ Briefly, constructed DGEList object (DGEList function), design matrix (model.design) and contrast matrix (makeContrasts) was defined, filtered low counts (filterByExpr), and trimmed mean of M-values (TMM) normalization (calcNormFactors) was done. Later, log_2_ counts per million reads (logCPM) transformation, model fitting and differential expression analysis was performed using limma-voom (voomWithQualityWeights, lmFit, contrasts.fit), empirical Bayes moderated t-statistics (eBayes) and TREAT method (treat, decideTests, topTreat) considering log fold-change (logFC) (0.5) threshold. Genes with the logFC (∼0.5) and FDR adjusted p-value cutoff (≤0.01) were considered as differentially expressed genes (DEGs). Gene ontology (GO) and pathway enrichment analysis was accomplished implementing ShinyGO (v0.77). ^93^ DEGs volcano plots were depicted utilizing EnhancedVolcano (v1.16.0) package. Standard GSEA was performed using GenePattern against the set of genes measured by Howard *et al.* to be downregulated (n = 191; log_2_FC < -0.5, p < 0.01) by C16 treatment in MV4:11 cells.^33, 65, 94^

### Extreme Limiting Dilution Assay (ELDA)

L0 and DI318 CSCs were split with Accutase® as single cells into NBMc. 200 µL of CSCs were seeded into each well of a 96-well TC treated plate (n = 10 well per density) at 50, 25, 13, 6, 3, and 1.5 cells/well. One full plate was treated with either DMSO (0.2% v:v) or the indicated doses of **C3TD078**. Spheroids were allowed to grow for 14 days at 37 °C/ 5% CO_2_ followed by manual classification of each well for the binary presence or absence of sphere(s) (independently assessed by two scientists). The stem-cell frequency was quantified and treatments were compared to DMSO as described by Hu and Smyth.^95^

### *In Vitro* Tier 1 DMPK

All *in vitro* DMPK screening assays were performed at Q2 Solutions, Inc. (Indianapolis, IN, USA). ***Microsomal Intrinsic Clearance:*** Test compounds (1 µM) were incubated with 1 mg/mL of pooled liver microsomes in phosphate buffer at 37 °C. Samples were taken at 0, 5, 10, 15, 30 and 45 min and analyzed by LC-MS. Testosterone was used as the positive control. Two replicates were obtained and average CL_int_ values are reported. ***Plasma Protein and Brain Homogenate Binding:*** Plasma protein binding assays in human and rat plasma were performed using equilibrium dialysis method and a fast gradient elution LC-MS/MS to estimate the percent of compound which binds to protein over a 4.5 hour incubation period at 37 °C, while undergoing orbital shaking. Assay was performed using a HT dialysis micro equilibrium device using dialysis membrane strips. A time 0 sample was taken after protein matrix is created and samples were taken from both the protein side and buffer side of the membrane after incubation and the parent compound was quantified. Brain homogenate binding was performed similarly as above using human brain homogenate, equilibrium dialysis, and LC-MS/MS methods. Fraction unbound is calculated by dividing the concentration of the buffer side by the concentration of the matrix side (plasma or brain). ***MDCK Passive Permeability:*** MDCK (Madin-Darby canine kidney) assay plates were seeded 3-4 days prior to running the assay. PET 24- well plates were seeded at a cell density of 0.875 x 10^5^ cells/well in a 250 µL apical well volume (3.52 x 10^5^ cells/mL) with a 1.0 mL volume of growth medium to the 24-well basolateral wells. The basolateral wells were rinsed once and the apical wells were rinsed twice with HEPES Buffer Saline (HBSS), and fresh HBSS was added to the assay plate in a 250 µL apical well volume and a 1.0 mL basolateral well volume. The apical chamber was incubated with Pgp inhibitor LSN335984 for 36 minutes at 37 °C and then replaced with test article and incubated for a 5 minute and 60 minute interval. Acetonitrile was added to each well of the basal chamber of the assay plate, which is mixed with the existing buffer and transferred to the LCMS plate for analysis using a SCIEX API 4000/5000 instrument. Dexamethasone and Atenolol were utilized as controls. Compound transport was measured in the absorptive direction and expressed as the percentage transport over the incubation period.

### *In situ* Brain Perfusion Technique

To evaluate the permeability across the blood-brain barrier of our novel WDR5 inhibitors in the absence of hepatic clearance we utilized the *in situ* brain perfusion technique. Mice were anesthetized via intraperitoneal injection of 100 mg/kg ketamine and 80 mg/kg xylazine. The peritoneal cavity was exposed, the descending aorta was clamped, and the right atrium was cut to prevent venous return to the heart. A 28G butterfly winged infusion catheter (Terumo Medical Care Solutions, Somerset, NJ, USA) was inserted into the left ventricle and mice were perfused with a modified physiological buffer containing 10 μM WDR5 inhibitor.^96^ Perfusion fluid pumped through the left cardiac ventricle and maintained at a constant rate 2.5 mL/min using a Pump 11 Elite Syringe Pump (704500, Harvard Apparatus, Holliston, MA, USA). To determine brain uptake of WDR5 inhibitors, mice were perfused for 120 s. The experiment was concluded by decapitation of the mouse. Following termination, the brain was rapidly removed and placed on ice. Inhibitor concentration in both brain and blood was determined by liquid chromatography tandem mass spectroscopy (Q2 Solutions, Indianapolis, IN, USA). The unidirectional uptake transfer constant (K_in_) was calculated by the following relationship:^97^

Where Q_br_ is the concentration of WDR5 inhibitor in brain (ng/g) corrected for residual inhibitor remaining in cerebrovasculature at the end of the perfusion time frame, C_pf_ is the perfusion fluid concentration of WDR5 inhibitor (ng/mL), T is the time length of the perfusion experiment, and V_0_ is the vascular volume of the WDR5 inhibitor (mL/g). For these experiments, an average value of 0.012 mL/g was used for vascular volume. ^98, 99^

### *In Vivo* Intraperitoneal Plasma:Brain Level (PBL) Studies

*In vivo* PBL studies for **C16** and **CCF343** were performed by the Rat Metabolic Physiology Core (RMPC) at the Vanderbilt University Medical Center (VUMC) Metabolic Physiology Shared Resource Core (MPSR) supported in part by Diabetes Research Training Center (DRTC) NIH grant DK020593. The studies were performed according to guidelines approved by the Institutional Animal Care and Use Committee (IACUC) of VUMC following the guidance of the Association for Assessment and Accreditation of Laboratory Animal Care (AAALAC). Male CD-1 mice (Charles River Laboratories, Wilmington, MA) were overnight fasted on the evening prior to study. On the morning of study mice were weighed and allowed to acclimate to the room for at least 30 minutes prior to dosing. Food was returned 3 hr after intraperitoneal dosing. At time zero IP of test article was given (20% β-HPCD, 1-10 mg/mL with addition of 1N HCl per molar equivalent of test article, N=8 mice, two per timepoint). At 0.5, 1, 3, and 6 h post-dose, mice were placed into a plane of anesthesia using Isoflurane. A terminal blood sample was collected via cardiac puncture followed by immediate euthanasia and brain collection. Brain was washed with cold PBS or Saline, blotted dry on a piece of gauze, weighed, and flash frozen in liquid nitrogen. Whole blood was centrifuged at 5000 x *g* for 5 minutes and plasma was removed into a fresh tube for storage. All samples were stored at -80 °C until shipment on dry ice for bioanalysis at Q2 Solutions. For **C3TD078**, PBL studies were conducted at Medicilon Inc. DMPK (CRO, Shanghai) using a similar protocol as above using modified timepoints (5 min, 15 min, 30 min, 1, 3, and 8 h), male ICR mice, and a formulation of 10 mg/mL of test article in 5% NMP + 10% Solutol + 85% (20% HP-β- cyclodextrin). The blood was taken via submandibular vein or other suitable vein, 0.03 mL/time point. Samples were placed in tubes containing K2-EDTA and stored on ice until centrifuged. The blood samples were centrifuged at 6800 x *g* for 6 minutes at 2-8°C within 1h after collection and stored frozen at -80°C. After blood sample collection, the brain was removed, rinsed with saline, dried with filter paper, placed into labeled EP tubes (1 tube/tissue/animal/time point, and frozen at -80°C prior to bioanalysis. The analytical results were validated using a calibration curve to lower limit of quantification of 2 ng/mL for parent analyte with tolbutamide as an internal standard.

### Short-Term Flank DI318 Xenograft

NOD-scid (NSG) mice were inoculated subcutaneously in the right flank with 2 × 10⁶ DI318 GSCs suspended in 100 μL of PBS:Matrigel (1:1). Over the next 6-7 days tumors developed to the point where a small mass was visual. A digital caliper measurement (length and width) were taken to get a baseline tumor size. Mice were randomized into treatment groups (vehicle or **C3TD078**), ensuring even distribution of biological sex across groups. Compounds were formulated in 20% HP β-cyclodextrin (BCD) and administered once daily via IP injection at 30 mg/kg for 7 consecutive days. Ellipsoid tumor volume was calculated as previously described and compared using a t-test.^100^

### Long-Term Intracranial L0 Xenograft

Intracranial xenografts were conducted as previously described. ^100, 101^ Briefly, L0 cells were intracranial injected into 6-12 week old NSG mice. Once fully anesthetized, mice were secured into a stereotaxic apparatus and 20,000 L0 Cells were injected into the left hemisphere 0.5 mm rostral and 1.8 mm lateral to the bregma with 3.5 mm depth from the scalp. One week post tumor implantation, animals were treated with for 80 days with 30 mg/kg QD IP **C3TD343** formulated in 20% BCD. Animals were monitored over time for the presentation of neurological and behavioral symptoms at which time the animals were humanely euthanized. Survival analysis was completed using PRISM 9.1.2 and comparison were made using Log-Rank (Mantel-Cox) test.

### Compound Synthesis and Characterization

Dihydroisoquinoline-based WDR5 inhibitors **C16** was synthesized and characterized as described previously.^37^ Triazole-based WDR5 inhibitors were synthesized according to **Scheme 1**. Details regarding synthetic method development will be reported elsewhere in due course. Briefly, starting from readily available methyl 3,5-dibromo-4- methylbenzoate **1**, the dibromo-1,5-disubstitued-1,2,3-tiazole key intermediate was achieved in six-steps. A novel one-pot tandem Suzuki Cross-Coupling and subsequent intramolecular C-H activation was utilized to form the 1,4,5-trisubstituted-1,2,3-triazole scaffold, the average yield for tricyclic formation was 50%. A subsequent three-step sequence involving deprotection, activation, and displacement yielded the fully functionalized target with the S2 warhead installed. An exemplary synthesis of **C16-TZ** is detailed below.

**Scheme 1.**
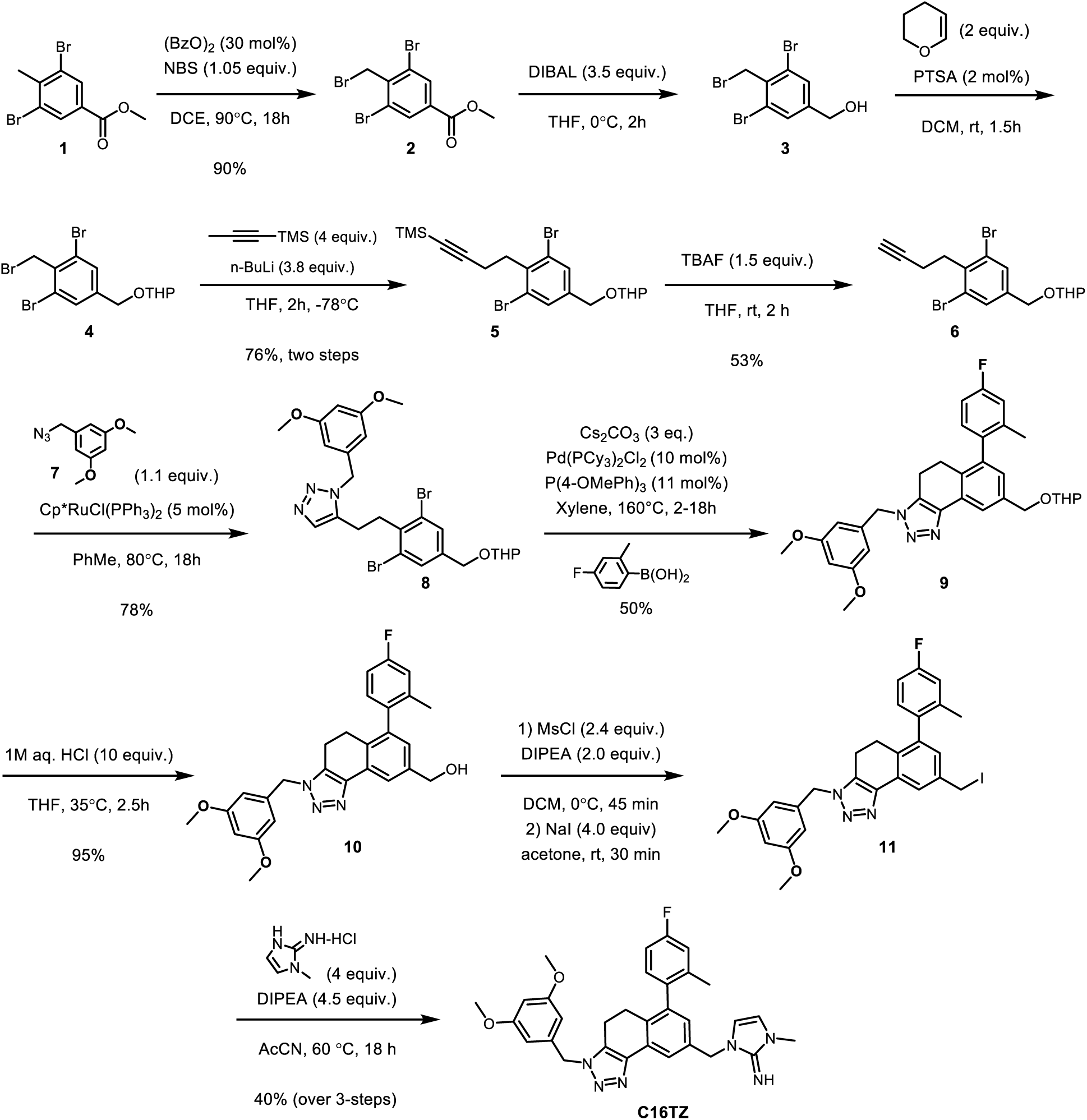
Synthesis of triazole-based WDR5 inhibitor **C16-TZ**.

1-((3-(3,5-Dimethoxybenzyl)-6-(1-ethyl-3-(trifluoromethyl)-1H-pyrazol-4-yl)-4,5-dihydro- 3H-naphtho[1,2-d][1,2,3]triazol-8-yl)methyl)-3-methyl-1,3-dihydro-2H-imidazol-2-imine (**C16-TZ**):

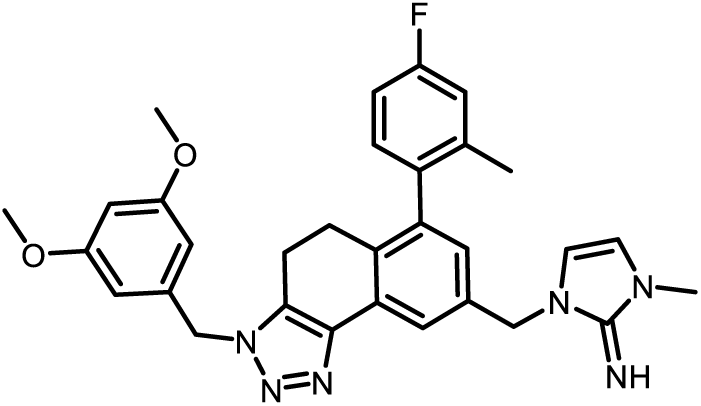

***Step 1.*** Preparation of methyl 3,5-dibromo-4-(bromomethyl)benzoate (**2**). To a 500 mL round bottom flask with a stir bar, the following were charged in the following order: methyl 3,5-dibromo-4-methyl-benzoate (**1**, 1.0 equiv., 24 g, 77.93 mmol), *N*-bromosuccinimide (1.05 equiv., 14.56 g, 81.8 mmol), benzoyl peroxide (0.3 equiv., 5.66 g, 23.3 mmol) and DCE (243.5 mL). The reaction mixture was heated to 90 °C and allowed to stir 18 h. The reaction mixture was cooled to room temperature and sat. NH_4_Cl (50 mL) added. The organic layer was separated, and the aq. layer was extracted with DCM (3 x 40 mL). The combined organic layers were washed with H_2_O (2 x 25 mL), sat. brine (50 mL), and dried over Na_2_SO_4_. The solids were filtered off, washed with DCM (2 x 25 mL) and the filtrate was concentrated under vacuum to give ∼27 grams of **2** as a white solid (90%). The crude solid was used without purification: ^1^H NMR (400 MHz, CDCl_3_) δ 8.19 (s, 2H), 4.82 (s, 2H), 3.93 (s, 3H).

***Step 2.*** Preparation of (3,5-dibromo-4-(bromomethyl)phenyl)methanol (**3**). To a 500 mL round bottom flask with a stir bar, **2** (methyl 3,5-dibromo-4-(bromomethyl)benzoate, 1.0 equiv., 10.5 g, 27.14 mmol) was dissolved with THF (86 mL) and the solution placed under argon atmosphere. The solution was cooled to 0 °C and 1M diisobutylaluminium hydride in THF (3.7 equiv., 100 mL) was added dropwise to the solution. The reaction mixture was stirred at 0 °C for 2.5 h. The reaction mixture was diluted with Et_2_O (100 mL), quenched at 0 °C by careful addition of 3M aq. NaOH (5 mL) followed by H_2_O (10 mL) dropwise (highly exothermic). The quenched solution was warmed to room temperature, stirred for 15 min, charged with MgSO_4_ (20 g) and then stirred for 15 min. The solids were filtered through a pad of Celite; the solids were mixed in EtOAc and filtered again. The collected filtrates were concentrated under vacuum to give a white solid upon further drying overnight (27.2 g) to give crude product **3** (>90% purity): ^1^H NMR (400 MHz, DMSO) δ 7.63 (s, 2H), 5.46 (s, 1H), 4.82 (s, 2H), 4.49 (s, 2H).

***Step 3.*** Preparation of 2-((3,5-dibromo-4-(bromomethyl)benzyl)oxy)tetrahydro-2*H*-pyran (**4**). To a 200 mL round bottom flask with a stir bar, the following were charged in the following order: **3** (3,5-dibromo-4-(bromomethyl)phenyl]methanol, 1.0 equiv., 9.74 g, 27.1 mmol), DCM (136 mL), 3,4-dihydro-2*H*-pyran (2.0 equiv., 4.95 mL, 54.3 mmol), and *p*- toluene sulfonic acid monohydrate (0.02 equiv., 103 mg, 0.54 mmol). The reaction mixture was stirred at room temperature for 1 hour. The reaction mixture was quenched with sat. NaHCO_3_ (10 mL) and vigorously stirred for 5 min. Sat. brine (15 mL) was then added and vigorously stirred for 5 mins. The organic layer was separated and dried over Na_2_SO_4_. The solids were filtered off, washed with DCM (2 x 25 mL) and the filtrate was concentrated under vacuum to give a crude yellow oil. The crude oil was purified by automated flash chromatography using a 330 g silica column and a gradient of 0-20% EtOAc in hexanes. The desired fractions were pooled and concentrated under vacuum to give the title product **4** as a clear yellow oil (9.14 g, 20.6 mmol, 76% two steps): ^1^H NMR (400 MHz, CDCl_3_) δ 7.56 (s, 2H), 4.93 (s, 2H), 4.72 – 4.67 (m, 2H), 3.86 (ddd, *J* = 11.3, 8.7, 3.4 Hz, 1H), 3.56 (ddd, *J* = 10.9, 5.4, 2.9 Hz, 1H), 1.92 – 1.54 (m, 7H).

***Step 4.*** Preparation of (4-(2,6-dibromo-4-(((tetrahydro-2*H*-pyran-2- yl)oxy)methyl)phenyl)but-1-yn-1-yl)trimethylsilane (**5**). To a 200 mL round bottom flask with a stir bar under Argon, THF (39.1 mL) and trimethyl(prop-1-ynyl)silane (4.0 equiv., 3.48 mL, 23.5 mmol) were charged. The solution was cooled to -78 °C, 2.5M *n*-BuLi in THF (3.8 equiv., 8.92 mL, 22.3 mmol) was charged dropwise; the reaction mixture was stirred for 2 h at -78 °C. A solution of **4** (2-[[3,5-dibromo-4- (bromomethyl)phenyl]methoxy]tetrahydropyran) from step 3 (1.0 equiv., 2.6 g, 5.87 mmol) in THF (19.57 mL) was charged dropwise; the reaction mixture was stirred for 2h at -78°C. At -78 °C, the reaction was quenched with MeOH (20 equiv. 4.75 mL) and the solution was allowed to warm. At 2-8 °C, EtOAc (60 mL), sat. NH_4_Cl (20 mL), H_2_O (10 mL) were charged and the mixture was warmed to room temperature. The organic layer was separated, and the aq. layer was extracted with EtOAc (30 mL). The combined organic layers were washed with sat. brine (20 mL), dried over Na_2_SO_4_, the solids filtered off, then washed with EtOAc (2 x 15 mL). The filtrate was then concentrated under vacuum to give a crude oil. The crude oil was purified by automated flash chromatography using an equilibrated 80 g silica column with 0-20% EtOAc in hexanes gradient. The desired fractions were pooled and concentrated under vacuum to give **5** as a clear yellow oil (2.11 g,4.45 mmol, 76% yield): ^1^H NMR (400 MHz, CDCl_3_) δ 7.50 (s, 2H), 4.72 – 4.63 (m, 2H), 4.39 (d, *J* = 12.5 Hz, 1H), 3.92 – 3.82 (m, 1H), 3.60 – 3.50 (m, 1H), 3.22 (t, *J* = 8.0 Hz, 2H), 2.51 (dd, *J* = 16.1, 8.0 Hz, 2H), 1.86 (qd, *J* = 11.6, 9.3, 2.6 Hz, 1H), 1.80 – 1.70 (m, 1H), 1.70 – 1.47 (m, 5H), 0.15 (s, 9H).

***Step 5.*** Preparation of 2-[(3,5-dibromo-4-but-3-ynyl-phenyl)methoxy]tetrahydropyran (**6**). To a 100-mL round bottom flask with a stir bar, intermediate **5** (4-[2,6-dibromo-4- (tetrahydropyran-2-yloxymethyl)phenyl]but-1-ynyl-trimethyl-silane, 1.0 equiv., 8 g, 16.9 mmol) was dissolved with THF (56 mL). A solution of TBAF (1M in THF, 1.5 equiv., 25.3 mL, 25.3 mmol) was charged at 23 °C. The reaction was stirred for 2 h at 23 °C. EtOAc (125 mL), sat. NH_4_Cl (40 mL) and H_2_O (20 mL) were charged and allowed to stirred for 10 min. The organic layer was separated, and the aq. layer was extracted with EtOAc (2 x 75 mL). The combined organic layers were washed with brine, dried over Na_2_SO_4_, the solids were filtered off and washed with EtOAc (2 x 20 mL). The filtrate was concentrated under vacuum to give a crude yellow oil. The crude oil was purified by automated flash chromatography using an equilibrated 330 g silica column and a 0-20% EtOAc in hexanes gradient. The desired fractions were pooled and concentrated under vacuum to give the desired product **6** as a clear yellow oil (5.5 g, 13.68 mmol, 81% yield): ^1^H NMR (400 MHz, CDCl3) δ 7.51 (s, 2H), 4.75 – 4.65 (m, 2H), 4.40 (d, *J* = 12.5 Hz, 1H), 3.92 – 3.83 (m, 1H), 3.61 – 3.51 (m, 1H), 3.31 – 3.20 (m, 1H), 2.56 (s, 1H), 2.47 (ddd, *J* = 9.8, 6.8, 2.6 Hz, 1H), 2.02 (t, *J* = 2.6 Hz, 1H), 1.97 – 1.56 (m, 7H).

***Step 6.*** Preparation of 5-(2,6-Dibromo-4-(((tetrahydro-2*H*-pyran-2- yl)oxy)methyl)phenethyl)-1-(3,5-dimethoxybenzyl)-1*H*-1,2,3-triazole (**8**). To a 100 mL round bottom flask with a stir bar, **6** (2-[(3,5-dibromo-4-but-3-ynyl- phenyl)methoxy]tetrahydropyran, 1.0 equiv., 5.g, 12.43mmol), 1-(azidomethyl)-3,5- dimethoxy-benzene (**7**, 1.2 equiv., 2.9 g, 14.9 mmol), toluene (51.8 mL), chloro(pentamethylcyclopentadienyl)bis(triphenylphosphine)ruthenium(II) (0.045 equiv., 445.6 mg, 0.56 mmol) were charged. The mixture was sparged with argon for 20 min.

The reaction was heated to 80 °C for 18 h. The reaction was cooled to room temp and the solids were filtered off and rinsed with EtOAc; the filtrate was concentrated under vacuum. The crude oil was diluted with minimal CH_2_Cl_2_, dried loaded and then purified by automated flash chromatography using a 120 g silica column (0-100% EtOAc:hexanes; product eluted at ∼70% EtOAc). The desired fractions were pooled and concentrated to give **8** as a viscous orange oil (5 g, 8.4 mmol, 68% yield): ^1^H NMR (400 MHz, CDCl_3_) δ 7.55 (s, 1H), 7.49 (s, 2H), 6.34 (t, *J* = 2.2 Hz, 1H), 6.27 (d, *J* = 2.3 Hz, 2H), 5.46 (s, 2H), 4.70 – 4.64 (m, 2H), 4.38 (d, *J* = 12.6 Hz, 1H), 3.85 (ddd, *J* = 11.2, 8.4, 3.0 Hz, 1H), 3.71 (d, *J* = 1.9 Hz, 6H), 3.58 – 3.50 (m, 1H), 3.21 – 3.13 (m, 2H), 2.83 – 2.75 (m, 2H), 1.89 – 1.47 (m, 6H). ^13^C NMR (101 MHz, CDCl_3_) δ 161.30, 140.49, 137.35, 137.26, 136.12, 133.10, 131.41, 124.96, 105.07, 100.11, 98.17, 66.93, 62.30, 55.45, 51.73, 35.82, 30.48, 25.40, 21.51, 19.32.

***Step 7 and 8.*** Preparation of (3-(3,5-dimethoxybenzyl)-6-(4-fluoro-2-methylphenyl)-4,5- dihydro-3H-naphtho[1,2-d][1,2,3]triazol-8-yl)methanol (**10**). To a 100 mL round bottom flask a stir bar, 5-[2-[2,6-dibromo-4-(tetrahydropyran-2-yloxymethyl)phenyl]ethyl]-1-[(3,5- dimethoxyphenyl)methyl]triazole (1.0 equiv., 1.0 g, 1.68 mmol), 4-fluoro-2-methylphenyl)boronic acid (2 equiv., 0.52 g, 3.36 mmol), tris(4-methoxyphenyl)phosphane (0.11 equiv., 65 mg, 0.18 mmol), dichloropalladium;tricyclohexylphosphane (0.10 equiv., 124 mg, 0.10 mmol), Cs_2_CO_3_ (4.0 equiv., 2.19 g, 6.7 mmol) and *m*-xylene (28 mL) were charged. The mixture was sparged with argon for 20 min. The reaction mixture was refluxed to 150 °C and allowed to stir vigorously (1500 rpm). The reaction was monitored by LC-M. After 3 h, the reaction was cooled to room temp and diluted with EtOAc (150 mL) and H2O (40 mL). The aqueous layer was extracted with EtOAc (3 x 50 mL). The combined organic layers dried over Na2SO4, the solids were filtered and rinsed with EtOAc. The filtrate and combined organic layers were concentrated and dried under vacuum. The crude oil was dissolved with THF (15 mL) and then 1M aq. HCl (10 equiv., 16.8 mL, 16.8 mmol) was charged. The reaction was heated to 35 °C for 2.5 h. The reaction mixture was cooled to room temp and diluted with EtOAc (35 mL) and sat. bicarb. was charged, dropwise (∼6 mL), until the pH was basic. The aq. layer was extracted with EtOAc (2 x 15 mL). The combined organic layer was washed with brine, dried over Na_2_SO_4_, the solids were filtered off and rinsed with EtOAc; the filtrate was concentrated under vacuum. The crude oil was dissolved with minimal DCM and then purified by automated flash chromatography (0-100% EtOAc/hexanes, product elutes at 100% EtOAc). The desired fractions were pooled and concentrated under vacuum. The resulting foam was dissolved in minimal THF and while stirring triturated with addition of hexanes (∼40 v/v% THF/hexanes) to afford a white solid. The solid was filtered and rinsed using 40% THF:hexanes and then dried under vacuum to afford the desired carbinol product as a while solid (350 mg, 45% yield): Purity ≥95% by LCMS (210, 254 nm)_;_ t_R_ = 1.69 min, *m/z* = 460.20 [M+H]^+^, ^1^H NMR (400 MHz, CDCl_3_) δ 7.99 (s, 1H), 7.35 (s, 1H), 7.17 (s, 1H), 6.39 (s, 1H), 6.35 (s, 2H), 5.46 (s, 2H), 4.74 (s, 2H), 4.26 (q, *J* = 7.4 Hz, 2H), 3.74 (s, 6H), 2.80 (t, *J* = 7.6 Hz, 2H), 2.68 (t, *J* = 7.4 Hz, 2H), 1.56 (t, *J* = 7.4 Hz, 3H). ^19^F NMR (376 MHz, CDCl_3_) δ -60.21.

***Step 9.*** Preparation of 3-(3,5-dimethoxybenzyl)-6-(4-fluoro-2-methylphenyl)-8- (iodomethyl)-4,5-dihydro-3H-naphtho[1,2-*d*][1,2,3]triazole. To a 20 mL vial with a stir bar was charged (3-(3,5-dimethoxybenzyl)-6-(4-fluoro-2-methylphenyl)-4,5-dihydro-3H- naphtho[1,2-d][1,2,3]triazol-8-yl)methanol (1.0 equiv., 230 mg, 0.5 mmol), CH_2_Cl_2_ (5 mL) and DIPEA (2.0 equiv., 0.17 mL, 1.0 mmol). The solution was cooled to 0°C, methanesulfonyl chloride (0.05 mL, 0.6 mmol) was added to the solution and stirred for 20 mins. Water (10 mL) was added to the reaction mixture at 0°C and the aq. layer was extracted with DCM (3 x 15 mL). The combined organic layers were dried over Na_2_SO_4_ and concentrated under vacuo. The crude oil, (3-(3,5-dimethoxybenzyl)-6-(4-fluoro-2- methylphenyl)-4,5-dihydro-3H-naphtho[1,2-*d*][1,2,3]triazol-8-yl)methyl methanesulfonate was dissolved with acetone (10 mL). To this solution, sodium iodide (4.0 equiv., 60 mg, 0.4 mmol) was added and allowed to stir at room temperature for 30 mins. Sat. brine (7 mL) was added to the reaction mixture and extracted with EtOAc (2 x 10 mL). The combined organic layers were dried over Na_2_SO_4_ and concentrated under vacuo. The crude was dissolved with minimal DCM:THF (1:1) and purified by automated flash chromatography (0-45% EtOAc/hexanes). The desired fractions were pooled and concentrated under vacuo, to give 61 mg the desired product as a yellow oil (21% overall yield, two steps): Purity >95% by LCMS (215, 254 nm); t_R_ = 2.45 min, *m/z* = 571.09 [M+H]^+^

***Step 10.*** Preparation of **C16-TZ** 1-((3-(3,5-Dimethoxybenzyl)-6-(1-ethyl-3-(trifluoromethyl)-1*H*-pyrazol-4-yl)-4,5-dihydro-3*H*-naphtho[1,2-d][1,2,3]triazol-8-yl)methyl)-3-methyl-1,3-dihydro-2*H*-imidazol-2-imine. To a 1-dram vial with a stir bar, 3-(3,5-dimethoxybenzyl)-6-(4-fluoro-2-methylphenyl)-8-(iodomethyl)-4,5-dihydro-3H-naphtho[1,2-d][1,2,3]triazole (1.0 equiv., 28.5 mg, 0.05 mmol), 3-methyl-1H-imidazol-2-imine hydrochloride (20 mg, 0.15 mmol, 3 equiv.), acetonitrile (0.30 mL) and DIPEA (0.035 mL, 0.20 mmol, 4 equiv.) were charged. The mixture was warmed to 60 °C and stirred for 18 h. The reaction mixture was cooled to room temp and the solids were filtered off. The filtrate was concentrated under vacuum. The crude oil was diluted by minimal DMSO and purified by reverse phase. The desired fractions were pooled and concentrated under positive nitrogen stream. The residue was diluted in EtOAc and washed with sat. K_2_CO_3_ to remove TFA. The organic layers were washed with brine, filtered from Na_2_SO_4_, and dried under vacuum to afford the title product as a white solid (13 mg, 48%): Purity ≥95% by LCMS (210, 254 nm)_;_ t_R_ = 2.048 min, *m/z* = 539.23 [M+H]^+^; ^1^H NMR (400 MHz, MeOD) δ 7.73 (d, *J* = 2.0 Hz, 1H), 7.12 – 7.00 (m, 2H), 7.00 – 6.91 (m, 2H), 6.70 – 6.56 (m, 2H), 6.46 – 6.40 (m, 1H), 6.37 (d, *J* = 2.2 Hz, 2H), 5.52 (s, 2H), 5.03 – 4.95 (m, 2H), 3.73 (s, 6H), 3.39 – 3.32 (m, 3H), 2.81 – 2.61 (m, 4H), 2.04 (s, 3H).

(6-Cyclopropyl-3-(4-fluoro-3-methoxybenzyl)-8-((2-methyl-1H-imidazol-1-yl)methyl)-4,5-dihydro-3H-naphtho[1,2-d][1,2,3]triazole (**C3TD343**):

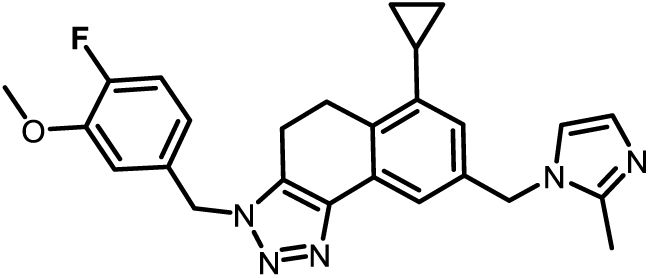

**C3TD343** was prepared following the procedures described in **C16-TZ**, substituting 4-(azidomethyl)-2-methoxy-1-fluoro-benzene (1.2 equiv., 652 mg, 3.6 mmol) in Intermediate **7** reacting with common intermediate dibromide **6** (1.2 g, 3 mmol) to afford the respective triazole intermediate **8** (1.5 g, 85% yield, 5-[2-[2,6-dibromo-4-(tetrahydropyran-2-yloxymethyl)phenyl]ethyl]-1-[(4-fluoro-3-methoxy-phenyl)methyl]triazole). Using 309 mg (0.53 mmol) of 5-[2-[2,6-dibromo-4-(tetrahydropyran-2-yloxymethyl)phenyl]ethyl]-1-[(4-fluoro-3-methoxy phenyl)methyl]triazole into Step 7 the two-step/one-pot CH activation ring closure and Suzuki reaction substituting cyclopropylboronic acid (1.1 equiv., 45 mg, 0.53 mmol) proceed smoothly. After flash chromatography purification (0-100% EtOAc/hexanes) 144 mg of the desired respective tricyclic core was isolated (58% yield). The subsequent deprotection, activation, and displacement (Step 10) steps substituting 2-methylimidazole as the S2 amine was performed on 0.7 mmol scale in the final step (4 equiv., 229 mg, 2.8 mmol). After purification using RP-HPLC the product fractions were pooled, concentrated under nitrogen stream and neutralized using NaOH. The free base product was extracted using DCM, washed with brine, filtered from Na_2_SO_4_, and dried under vacuum to afford 42 mg the title product as a white solid (15%): Purity ≥95% by LCMS (210, 254 nm); t_R_ = 1.87 min, *m/z* = 444.52 [M+H]^+^; ^1^H NMR (400 MHz, MeOD) δ 7.50 (s, 1H), 7.12 – 7.02 (m, 3H), 6.86 (s, 1H), 6.83 – 6.73 (m, 2H), 5.57 (s, 2H), 5.13 (s, 2H), 3.84 (s, 3H), 3.22 (t, *J* = 7.9 Hz, 2H), 2.90 (t, *J* = 7.9 Hz, 2H), 2.31 (s, 3H), 1.97 – 1.86 (m, 1H), 0.98 – 0.89 (m, 2H), 0.59 – 0.53 (m, 2H).

(3-(3,5-Dimethoxybenzyl)-8-((2-methyl-1*H*-imidazol-1-yl)methyl)-6-(1-methyl-3-(trifluoromethyl)-1*H*-pyrazol-4-yl)-4,5-dihydro-3*H*-naphtho[1,2-*d*][1,2,3]triazole (**C3TD078**):

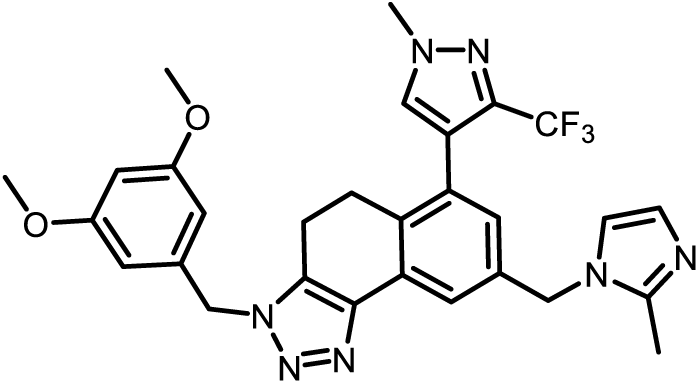

**C3TD078** was prepared following the procedures described in **C16-TZ**, starting from the common tricyclic S7 containing intermediate **8** the two-step/one-pot CH activation ring closure and Suzuki reaction (Step 7) was conducted substituting [1-methyl-3-(trifluoromethyl)pyrazol-4-yl]boronic acid (2 equiv., 391 mg, 2.0 mmol). Upon consumption of starting material as determined by LCMS the acid mediated deprotection was performed (Step 8). The crude was purified by automated flash chromatography purification (0-100% EtOAc/hexanes) to afford 190 mg of the desired respective tricyclic core carbinol (38% yield, Purity ≥95% by LCMS (210, 254 nm); t_R_ = 1.31 min, *m/z* = 505.52 [M+H]^+^). The subsequent activation and displacement (Steps 9-10) was performed using 2-methylimidazole as the S2 amine on 3 mmol scale in the final step (4 equiv., 1.3 g). After purification using RP-HPLC the product fractions were pooled, concentrated under nitrogen stream and neutralized using NaOH. The free base product was extracted using DCM, washed with brine, filtered from Na_2_SO_4_, and dried under vacuum to afford 310 mg the title product as a white solid (55%): Purity ≥95% by LCMS (210, 254 nm); t_R_ = 1.89 min, *m/z* = 564.23 [M+H]^+^; ^1^H NMR (400 MHz, MeOD) δ 7.71 (d, *J* = 13.2 Hz, 2H), 7.08 (s, 1H), 6.86 (s, 2H), 6.47 – 6.33 (m, 3H), 5.53 (s, 2H), 5.22 (s, 2H), 3.97 (s, 3H), 3.73 (s, 6H), 2.88 – 2.73 (m, 4H), 2.32 (s, 3H).

6-Cyclopropyl-3-[(4-methoxy-2-pyridyl)methyl]-8-[(2-methylimidazol-1-yl)methyl]-4,5-dihydrobenzo[e]benzotriazole (**C3TD424**):

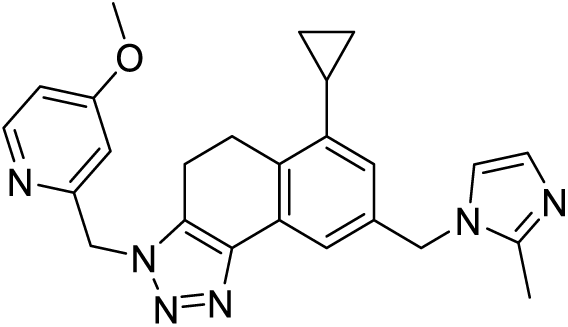

**C3TD424** was prepared following the procedures described in **C16-TZ**. Intermediate **6** was reacted in sequence with Steps 6-9, substituting 2-(azidomethyl)-4-methoxy-pyridine (1 equiv., 408.24 mg, 2.49 mmol) in Step 6 and substituting cyclopropylboronic acid (1.1 equiv., 136.16 mg, 1.59 mmol) in Step 7, and substituting 2-methylimidazole (2 equiv., 38.02 mg, 0.463 mmol) in the displacement Step 10, with addition of free basing the product TFA salt with aq. NaOH after RP-HPLC purification. Upon solvent removal under reduced pressure and further drying in vacuo the title product was obtained as a white solid (18 mg, 0.042 mmol, 18%): Purity >95% by LCMS (215, 254 nm); t_R_ = 1.431 min, *m/z* = 427.2251 [M+H]^+^; ^1^H NMR (400 MHz, CDCl3) δ 8.39 (d, J = 5.8 Hz, 1H), 7.70 (s, 1H), 7.01 (d, J = 2.1 Hz, 1H), 6.82 (s, 1H), 6.78 (s, 1H), 6.71 (s, 1H), 5.68 (s, 2H), 5.15 (d, J = 27.1 Hz, 2H), 3.84 (s, 3H), 3.25 (q, J = 7.9 Hz, 2H), 3.01 (t, J = 8.1 Hz, 2H), 2.80 (s, 3H), 1.25 (s, 1H), 1.03 – 0.93 (m, 2H), 0.56 (d, J = 5.4 Hz, 2H). ^13^C NMR (101 MHz, CDCl3) δ 166.76, 156.71, 150.88, 150.79, 140.35, 137.21, 136.58, 132.90, 131.34, 124.94, 109.35, 109.02, 108.01, 107.49, 98.07, 66.87, 62.24, 55.27, 53.26, 35.52, 30.42, 25.35, 21.51, 19.25.

## AUTHOR CONTRIBUTIONS

JAC designed and conducted the cellular experiments and wrote the manuscript. JAC, CMG, and AKG prepared figures. SAS conducted the *in vivo* perfusion experiments, advised on all pharmacokinetics, and conducted intracranial xenografts. SRM, SHH, JA, and SRS designed and synthesized the reported compounds. AKG and FC assisted with bioinformatic analyses. ARS, GB, JR, TN, NR, TR, and NSW assisted with the cellular experiments. PP, ER, AL, and AR performed protein purification, x-ray crystallography, and biochemical assays. ARS and GB conducted the flank xenografts. CMG solved the reported crystal structures. SAS, ARS, GB, CMG, NSW, JDL, and SRS provided editorial guidance for the manuscript, which was approved in its final form by all authors. JA, CGH, JDL, and SRS secured funding and provided supervision for the project.

## ACKNOWLEDGEMENT

This work was supported by the Falk Medical Research Trust Catalyst Awards Program and Cleveland Clinic Research. This research used resources of the Advanced Photon Source, a U.S. Department of Energy (DOE) Office of Science User Facility operated for the DOE Office of Science by Argonne National Laboratory under Contract No. DE-AC02-06CH11357. Use of the LS-CAT Sector 21 was supported by the Michigan Economic Development Corporation and the Michigan Technology Tri-Corridor (Grant 085P1000817).

