## Supplemental Information for "Development and Characterization of Triazole-Based WDR5 Inhibitors for the Treatment of Glioblastoma"

### Table of Contents


- Supplemental Figure 1: Plasma:blood level (PBL) study for **C16**
- Supplemental Figure 2: Data collection and refinement statistics for novel WDR5 co-crystal structures
- Supplemental Figure 3: X-ray crystal structures of WDR5 in complex with triazole based inhibitors with differing S2, S4, and S7 pieces
- Supplemental Figure 4: Permeability of **C16** and **C3TD343** in MDR1-MDCK cells
- Supplemental Figure 5: Volcano plots and biological processes GO enrichment for RNAseq study of **C16** and **C3TD879** in L0 and DI318 CSCs
- Supplemental Figure 6: Core set of WIN-site regulated genes in CSCs
- Supplemental Figure 7: Enriched GO terms and pathways in CSC RNAseq studies
- Supplemental Figure 8: CSC viability assays with **C16** and **C3TD879** in combination with radiation
- Supplemental Figure 9: Expression of *PYGB* in CSCs as assessed by RT-qPCR
- Supplemental Figure 10: L0 CETSA washout for **C16**
- Supplemental Figure 11: Chemical structure and *in vitro* profile of **C3TD424**
- Supplemental Figure 12: Phospholipidosis assay in A549 cells with **C16** and **C3TD879**
- Supplemental Figure 13: Cross-titration experiments with **C16** and **C3TD078** in combination with rationally selected anti-cancer drugs including Venetoclax
- Supplemental Figure 14: Protein expression of WDR5, ATAD2, and p53 in CSCs
- Supplemental Figure 15: Cross-titration experiments with **C3TD078** and the ATAD2 bromodomain inhibitor **GSK-8814** in five different CSC models.

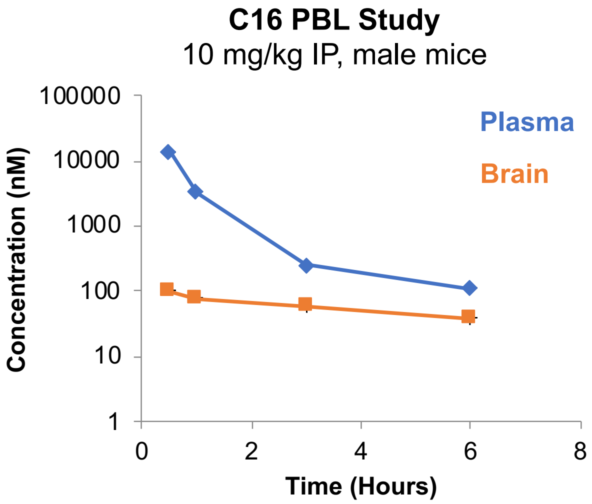


| | Plasma | Brain |
|---|---|---|
| AUC (nM*hrs) | 11,900 | 357 |
| AUC Interval (hrs) | 6 | 6 |
| $C_{max}$ (nM) | 14,200 | 102 |
| $T_{max}$ (hrs) | 0.5 | 0.5 |
| **P:B Ratio (AUC)** | 0.03 | |

**Supplemental Figure 1.** Snapshot plasma:blood level (PBL) study for **C16** dosed at 10 mg/kg IP in male mice. **C16** was formulated in 20% beta-cyclodextrin at 2 mg/mL. Raw concentration time-course (left, mean values from n = 2 mice) and extracted values (right) are shown.

| PDB ID | 9NCW | 9NCV | 9NCT |
|---|---|---|---|
| **Beamline** | APS LS-CAT 21-ID-F | APS LS-CAT 21-ID-F | APS LS-CAT 21-ID-F |
| **Wavelength (A)** | 0.97872 | 0.97872 | 0.97872 |
| **Resolution range** | 38.23 - 1.58 (1.6 - 1.58) | 45.5 - 1.583 (1.6 - 1.58) | 44.67 - 2.106 (2.16 - 2.11) |
| **Space group** | P 1 | P 1 | P 1 21 1 |
| **Unit cell** | 46.53 68.69 93.01 91.29 103.26 90.38 | 46.76 68.83 94.38 89.40 76.65 89.97 | 69.00 46.88 93.328 90.00 113.52 90.00 |
| **Total reflections** | 558098 | 538443 | 204477 |
| **Unique reflections** | 144679 (3473) | 148776 (4019) | 30647 (1951) |
| **Multiplicity** | 3.8 (3.3) | 3.6 (2.9) | 6.7 (6.0) |
| **Completeness (%)** | 94.95 (67.20) | 95.28 (76.15) | 95.45 (86.25) |
| **Mean I/sigma(I)** | 11.03 (1.06) | 14.52 (1.76) | 19.96 (3.26) |
| **Wilson B-factor** | 14.53 | 14.22 | 25.78 |
| **R-merge** | 0.101 (0.577) | 0.079 (0.305) | 0.117 (0.526) |
| **R-meas** | 0.125 (0.690) | 0.101 (0.438) | 0.138 (0.552) |
| **R-pim** | 0.064 (0.372) | 0.053 (0.246) | 0.053 (0.221) |
| **CC1/2** | 0.981 (0.836) | 0.992 (0.923) | 0.993 (0.862) |
| **CC*** | 0.995 (0.954) | 0.998 (0.980) | 0.998 (0.962) |
| **Reflections used in refinement** | 144679 (3473) | 148776 (4019) | 30647 (1951) |
| **Reflections used for R-free** | 7045 (182) | 7593 (168) | 1993 (125) |
| **R-work** | 0.2491 (0.4321) | 0.2528 (0.2870) | 0.1754 (0.2088) |
| **R-free** | 0.2753 (0.4238) | 0.2873 (0.3155) | 0.2282 (0.2751) |
| **Number of non-hydrogen atoms** | 10305 | 10456 | 4873 |
| **macromolecules** | 9074 | 9067 | 4539 |
| **ligands** | 80 | 160 | 68 |
| **solvent** | 1151 | 1229 | 266 |
| **Protein residues** | 1184 | 1184 | 592 |
| **RMS(bonds)** | 0.006 | 0.055 | 0.006 |
| **RMS(angles)** | 0.83 | 2.67 | 0.86 |
| **Ramachandran favored (%)** | 96.23 | 97.00 | 95.72 |
| **Ramachandran allowed (%)** | 3.77 | 3.00 | 4.28 |
| **Ramachandran outliers (%)** | 0 | 0 | 0 |
| **Rotamer outliers (%)** | 0.1 | 0.2 | 0.4 |
| **Clashscore** | 2.73 | 2.44 | 3.99 |
| **Average B-factor** | 16.97 | 16.08 | 27.01 |
| **macromolecules** | 16 | 15.09 | 26.8 |
| **ligands** | 19.44 | 16.61 | 29.06 |
| **solvent** | 24.43 | 23.34 | 30.06 |

**Supplemental Figure 2.** Table 1 data collection and refinement statistics for the three WDR5 co-crystal structures reported herein. One crystal was used for each structure. Values in parentheses unless stated represent data in the highest-resolution shell. RMS = root mean square.

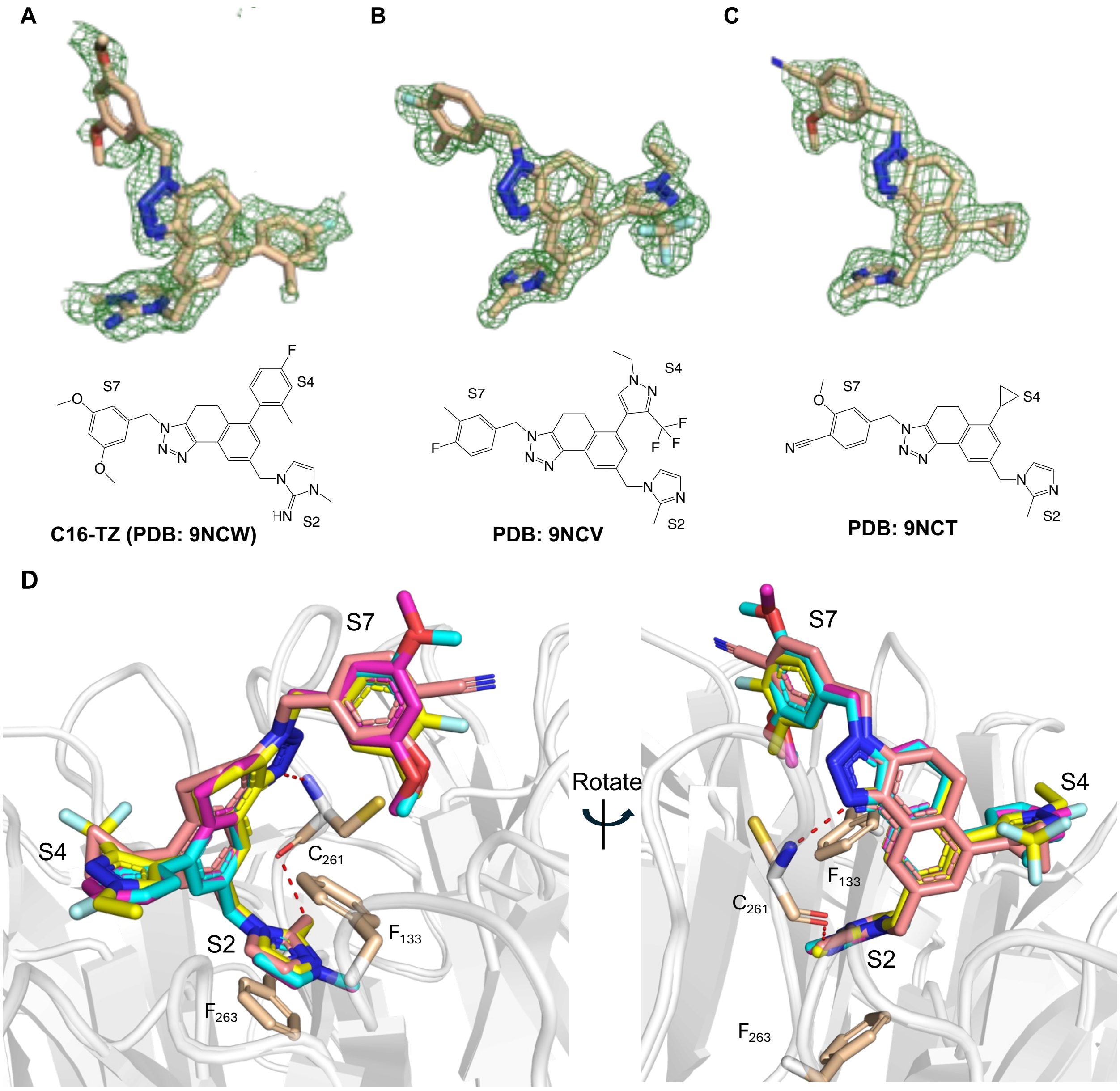


**Supplemental Figure 3.** Additional x-ray crystal structures of WDR5 in complex with triazole based inhibitors with differing S2, S4, and S7 pieces. Ligand Polder OMIT maps contoured to $\sigma$ = 3.0 and shown for Chain A ligands **(A) C16-TZ**, PDB: 9NCW), **(B)** PDB: 9NCV and (**C**) PDB: 9NCT. **(D)** Alignment of ligands highlights a conserved binding pose (**C16-TZ** – blue sticks, 9NCV – yellow sticks, 9NCT– pink sticks).

| Test Article | Direction | Recovery (%) | $P_{app}$ (10-6 cm/s) | | | Efflux Ratio | Brain Penetration Classification |
|---|---|---|---|---|---|---|---|
| | | | R1 | R2 | AVG | | |
| **C16** | A-to-B | 85.6 | 0.556 | 0.875 | 0.715 | 2.38 | Low |
| | B-to-A | 87.3 | 1.49 | 1.91 | 1.70 | | |
| **CCF343** | A-to-B | 71.2 | 0.131 | 0.0949 | 0.113 | 383 | Low |
| | B-to-A | 76.8 | 40.2 | 46.3 | 43.3 | | |

**Supplemental Figure 4.** Blood-brain barrier (BBB) penetration potential using MDR1-MDCK cell monolayers. All P-glycoprotein (MDR1) permeability and efflux studies were conducted at Absorption Systems (parent company Pharmaron). In brief, MDR1-MDCK monolayers were grown to confluence on collagen-coated, microporous membranes in 12-well assay plates. The permeability assay buffer was Hanks' balanced salt solution containing 10 mM HEPES and 15 mM glucose at a pH of 7.4. The buffer in the receiver chamber also contained 1% bovine serum albumin. The dosing solution concentration was 5 µM of test article in the assay buffer. Cell monolayers were dosed on the apical side (A-to-B) or basolateral side (B-to-A) and incubated at 37°C with 5% CO2 in a humidified incubator. Samples were taken from the donor and receiver chambers after 120 minutes. Each determination was performed in duplicate. The flux of lucifer yellow was also measured post-experimentally for each monolayer to ensure no damage was inflicted to the cell monolayers during the flux period. All samples were assayed by LC-MS/MS using electrospray ionization. Efflux ratio (ER) is defined as Papp (B-to-A) / Papp (A-to-B).

Brain Penetration Potential Classification:
$P_{app}$ (A-to-B) ≥ 3.0 ($10^{-6}$ cm/s) and ER < 3.0: **High**
$P_{app}$ (A-to-B) ≥ 3.0 ($10^{-6}$ cm/s) and 10 > ER ≥ 3.0: **Moderate**
$P_{app}$ (A-to-B) ≥ 3.0 ($10^{-6}$ cm/s) and ER ≥ 10, or Papp (A-to-B) < 3.0 ($10^{-6}$ cm/s): **Low**

**A**
**L0 CSCs, 200 nM C3TD078 vs. DMSO**

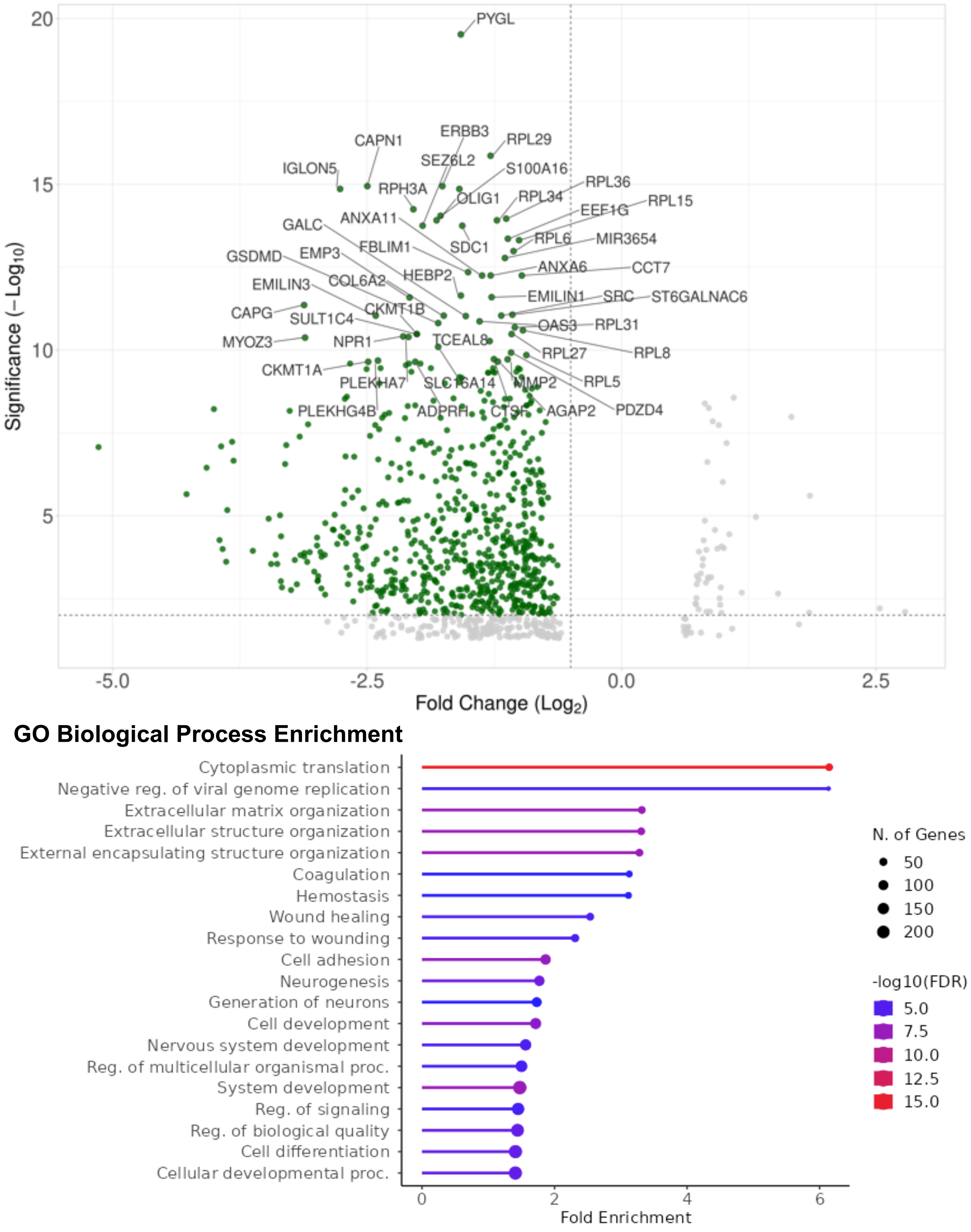

**B**

**DI318 CSCs, 200 nM C3TD078 vs. DMSO**

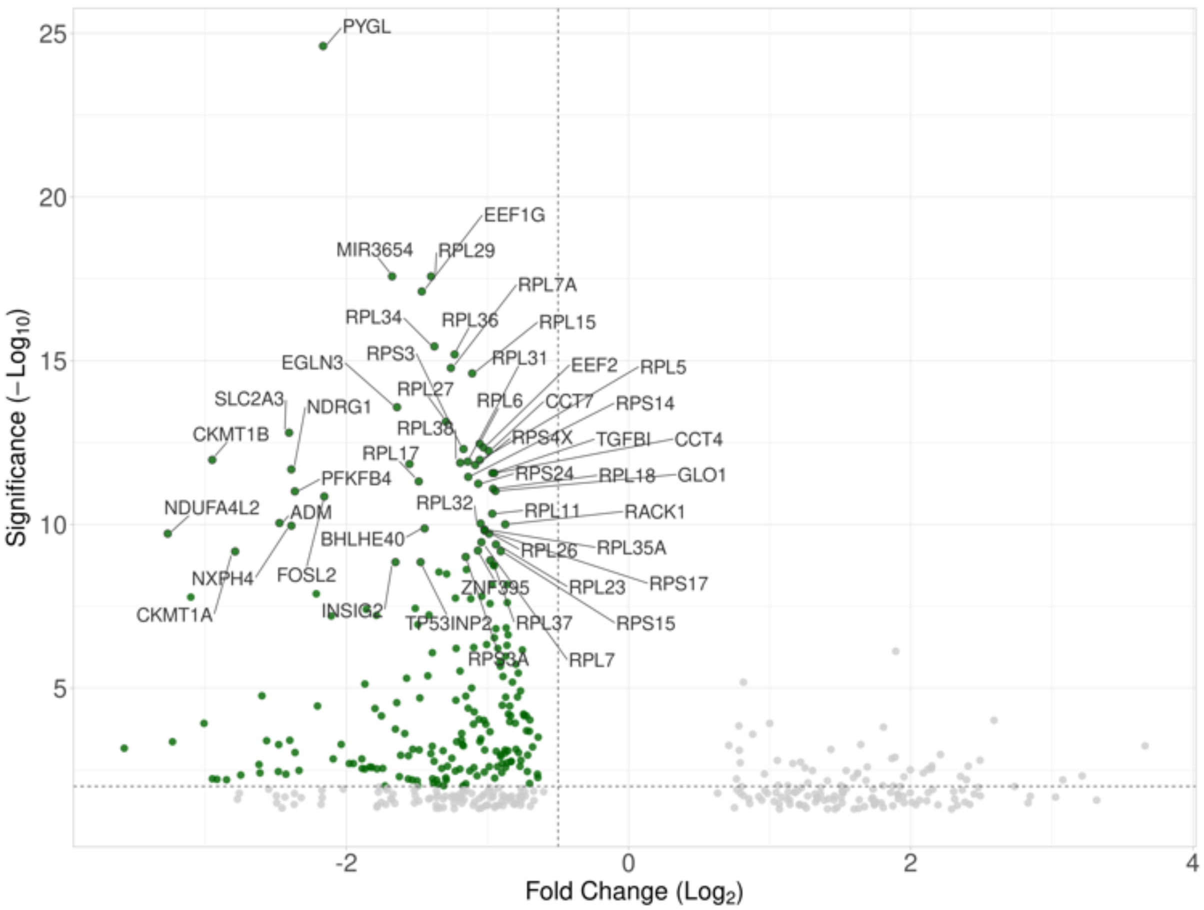


**GO Biological Process Enrichment**

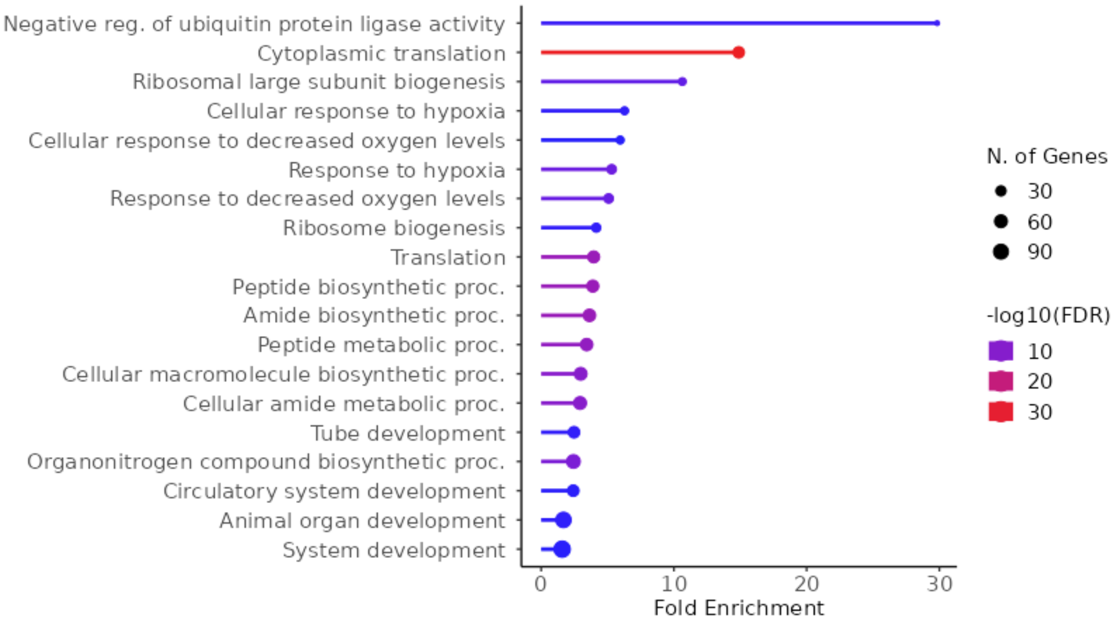

**C**
**L0 CSCs, 200 nM C16 vs. DMSO**

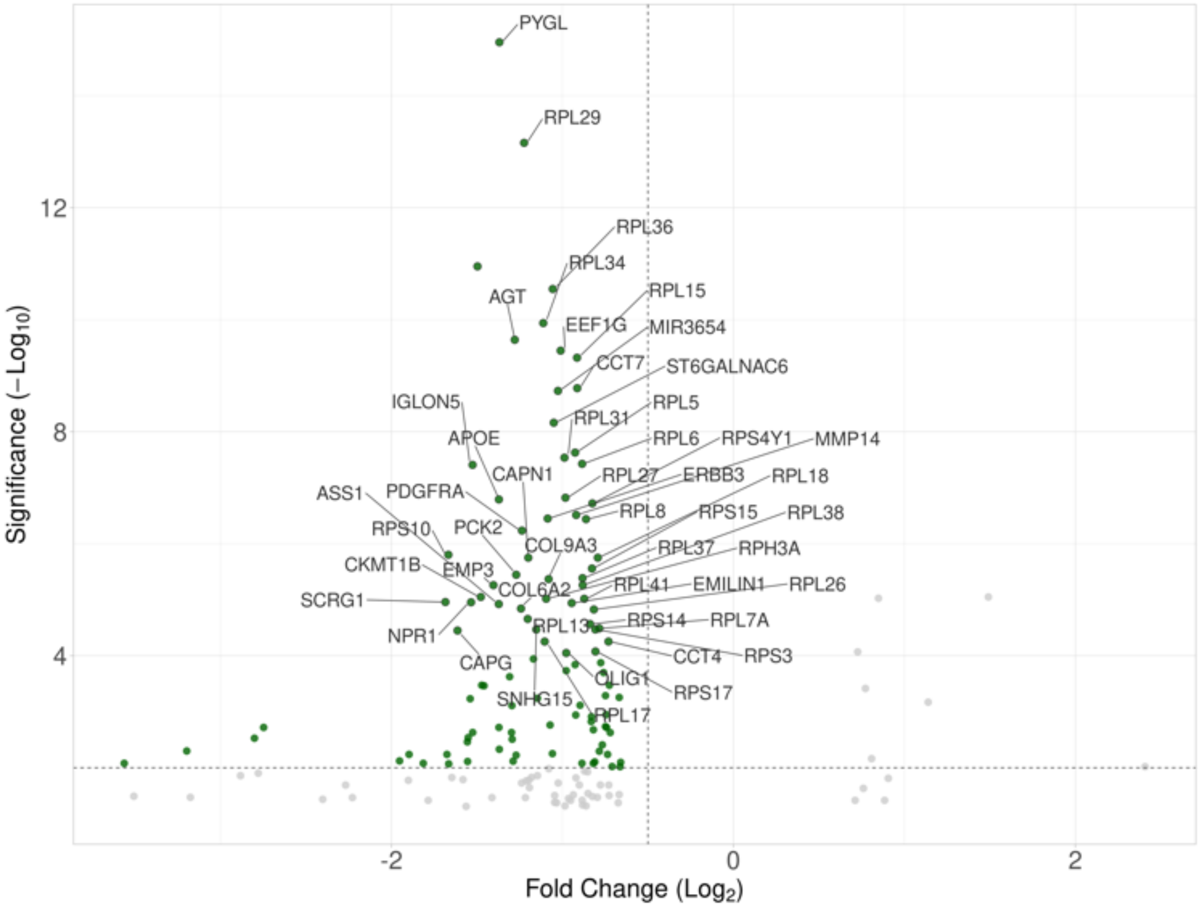


**GO Biological Process Enrichment**

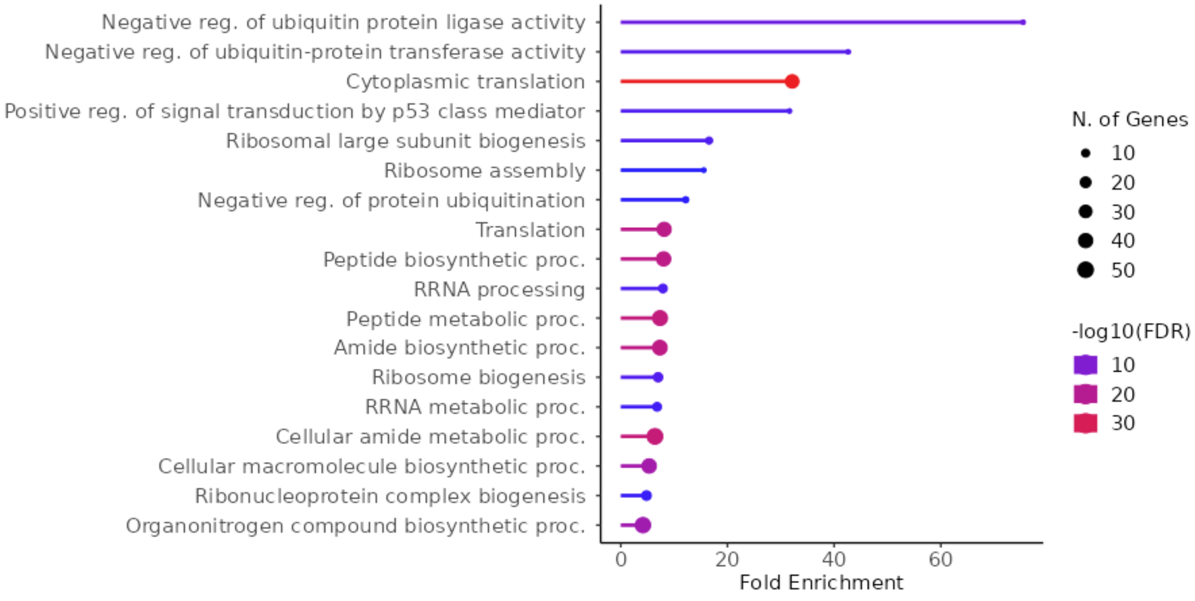

**D**

**DI318 CSCs, 200 nM C3TD078 vs. DMSO**

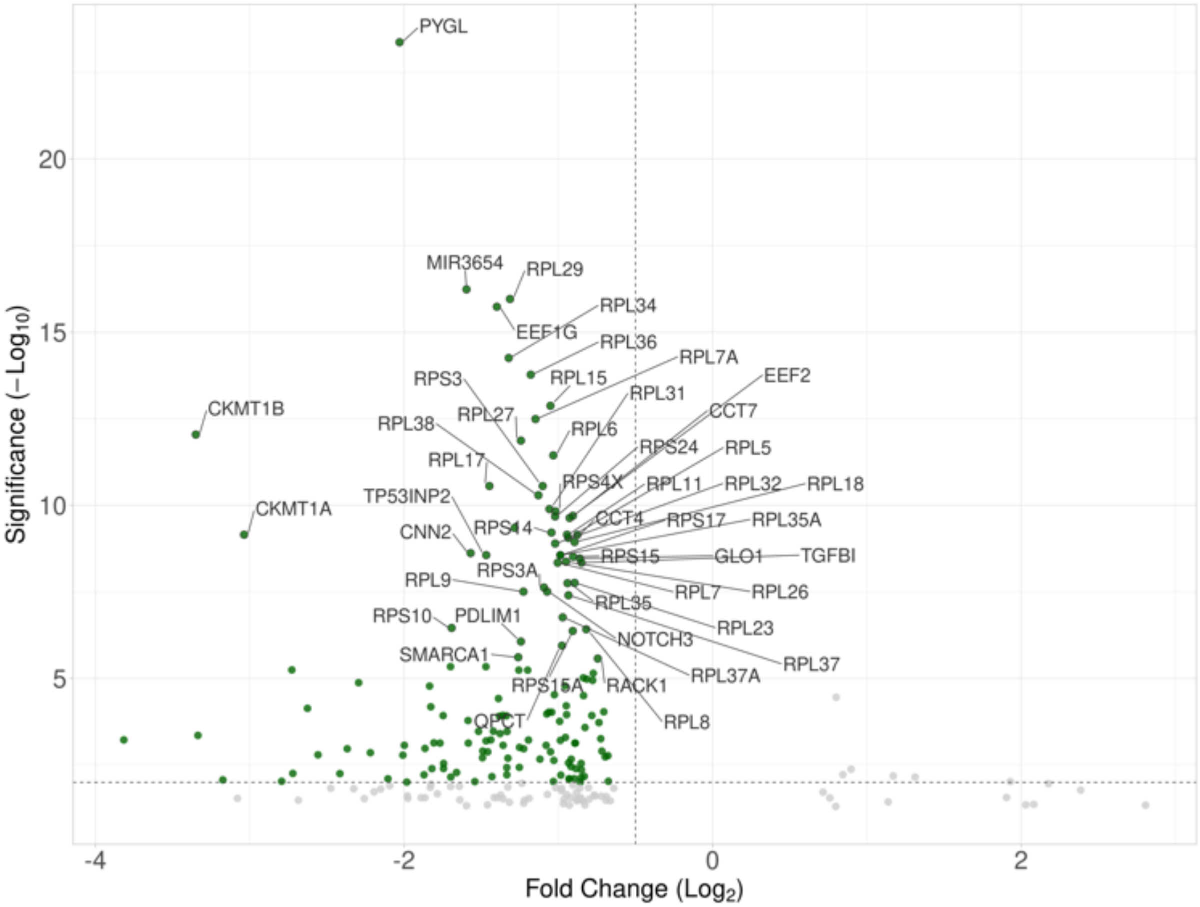


### GO Biological Process Enrichment

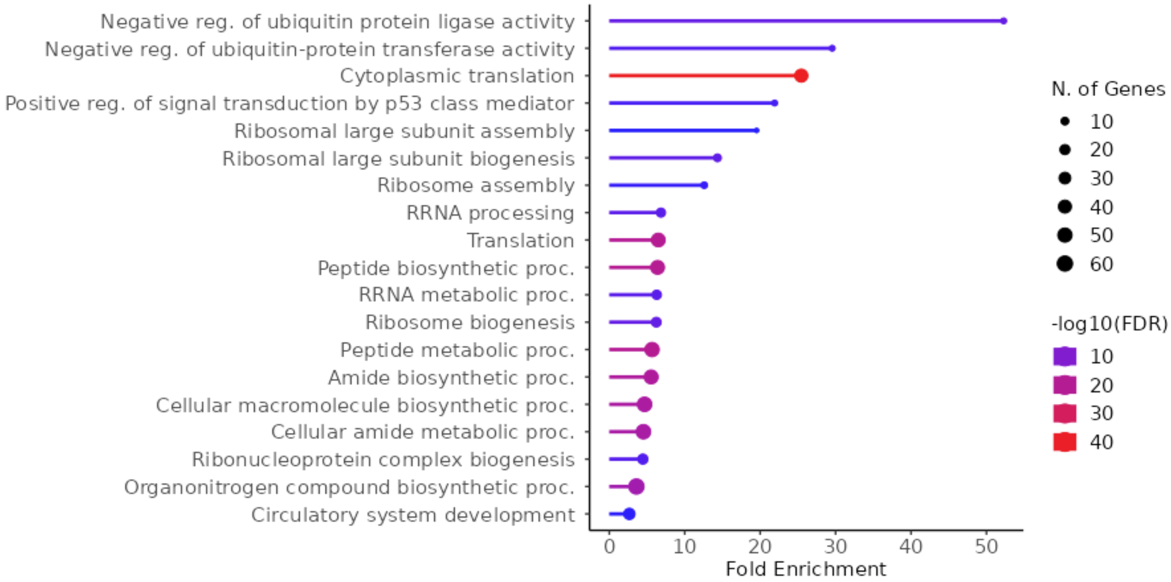

**Supplemental Figure 5.** Volcano plots (differentially downregulated genes in green, top 50 annotated) and biological processes GO enrichment for **C3TD879** in **(A)** L0 and **(B)** DI318 CSCs**,** as well as **C16** in **(C)** L0 and **(D)** DI318 CSCs.

- PYGL
- RPL29
- SDC1
- RPL34
- RPL36
- EEF1G
- RPL15
- RPL6
- MIR3654
- EMILIN3
- CCT7
- CKMT1B
- CKMT1A
- RPL31
- RPL8
- RPL27
- RPL5
- RPS10
- RPL41
- RPL38
- ELF4
- RPS17
- RPS15
- RPL9
- RPL26
- RPS24
- RPS14
- RPL37
- CCT4
- RPL17
- GLO1
- RPL32
- RPS4X
- RPS3A
- RPL7
- RPL23
- RPL7A
- RPS18
- RPL35A
- RPL18
- RPL35
- RPS3
- RPS15A
- RPL11

**Supplemental Figure 6.** Core set of WIN-site regulated genes in CSCs. These genes were downregulated ($p < 0.01$; $\log_2 FC < -0.5$) by both **C3TD078** and **C16** in both L0 and DI318 CSC models as measured by bulk RNAseq.

**A**

| GO TERM | L0 CSCs (FDR) | | DI318 CSCs (FDR) | |
|---|---|---|---|---|
| | C16 | CCF078 | C16 | CCF078 |
| GO:0002181 **Cytoplasmic translation** | $7.4 \times 10^{-40}$ | $2.6 \times 10^{-16}$ | $1.6 \times 10^{-41}$ | $1.5 \times 10^{-32}$ |
| GO:0042254 **Ribosome biogenesis** | $3.2 \times 10^{-6}$ | *n.s.* | $2.7 \times 10^{-7}$ | $1.4 \times 10^{-7}$ |
| GO:0022008 **Neurogenesis** | *n.s.* | $1.7 \times 10^{-6}$ | *n.s.* | *n.s.* |
| GO:0001666 **Response to hypoxia** | *n.s.* | *n.s.* | *n.s.* | $1.7 \times 10^{-8}$ |

**B**

| Reactome Pathway | Entities Found | Entities Total | FDR |
|---|---|---|---|
| **Translation** | 40 | 294 | $< 3.3 \times 10^{-16}$ |
| **rRNA Processing** | 39 | 203 | $< 3.3 \times 10^{-16}$ |
| **Peptide Chan Elongation** | 39 | 90 | $< 3.3 \times 10^{-16}$ |
| **rRNA Processing in the Nucleus and Cytosol** | 39 | 193 | $< 3.3 \times 10^{-16}$ |

**Supplemental Figure 7. (A)** Highlighted enriched GO terms (biologically processes) across cell lines and treatments. **(B)** Pathway enrichment analysis using Reactome for the "core" set of WIN-site dependent CSC genes highlighted in **Supplemental Figure 6**.

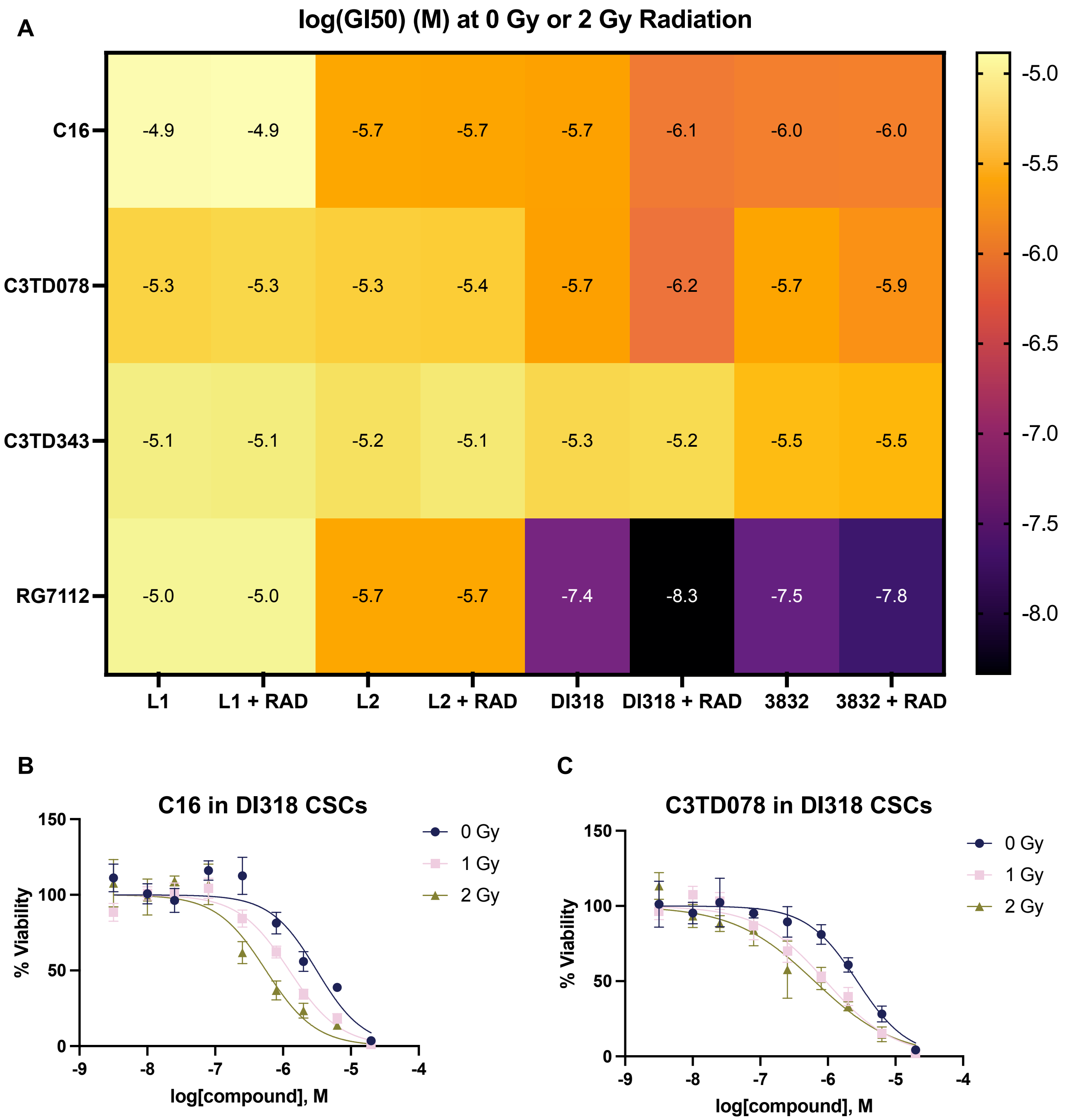


**Supplemental Figure 8**. **(A)** pGI50 (-log[GI50]] (M) values for L1, L2, DI318, and 3832 CSC models treated with a dose-response of **C16**, **C3TD078, C3TD343**, or the MDM2 antagonist (p53 activator) **RG7112** following either 0 Gy or 2 Gy irradiation. Mean values are presented from a single biological replicate conducted in technical triplicate. Follow-up experiments in just DI318 CSCs pre-treated for three days with either **(B) C16** and **(C) C3TD078** following irradiation with either 0, 1, or 2 Gy. Cell viability was quantified as described in the Materials and Methods four days post-irradiation (seven days total). The sensitivity of DI318 CSCs to WIN-site inhibitors was modestly enhanced by radiation.

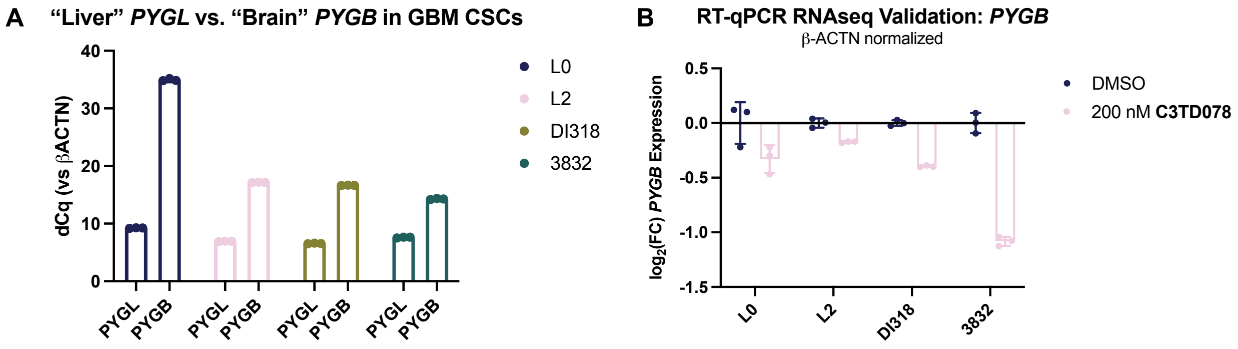


**Supplemental Figure 9**. **(A)** Expression (dCq vs ACTB) for the liver *PYGL* isoform and the brain *PYGB* isoform in L0, L2, DI318, and 3832 CSCs as quantified by RT-qPCR. The "liver" *PYGL* is expressed at a much higher level than the "brain" *PYGB*. **(B)** Unlike *PYGL,* the expression of *PYGB* is largely unaffected ($log_2FC < 1.0$) in L0, L2, DI318, and 3832 cells treated for 72 hours with 200 nM **C3TD078.**

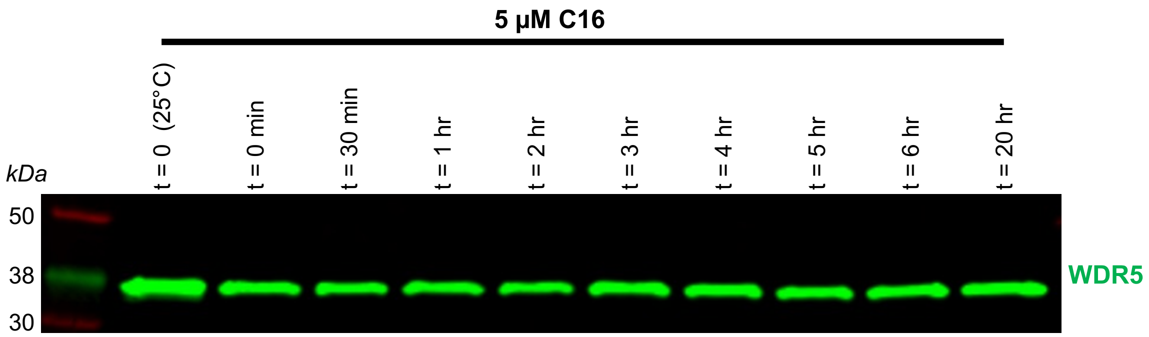


**Supplemental Figure 10**. L0 CSCs were treated for 2 hours with 5 µM **C16** prior to compound washout, and samples were prepared for CETSA WB at the indicated time points post-compound removal. The first sample was not heated and represents total cellular WDR5, while the other lanes were heated to 70°C to remove unbound WDR5 (see Materials and Methods). Like **C3TD078**, **C16** remains bound to WDR5 in L0 CSCs for at least 20 hours following washout.

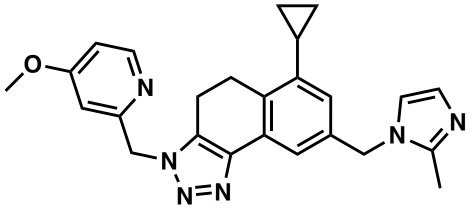

| | C3TD424 |
|---|---|
| MW (g/mol)<br>CLogP<br>TPSA (Å$^2$) | 426.52<br>2.3<br>65.2 |
| TR-FRET $K_i$ (nM) | 1.6 +/- .07 (n = 3) |
| CETSA $K_d$ (nM) | 500<br>(n = 1) |

**Supplemental Figure 11.** Chemical structure and *in vitro* profile of **C3TD424**, the weakly potent inhibitor related to **C3TD343**, that was used to validate the on-target nature of the transcriptional responses presented in **Figure 3**.

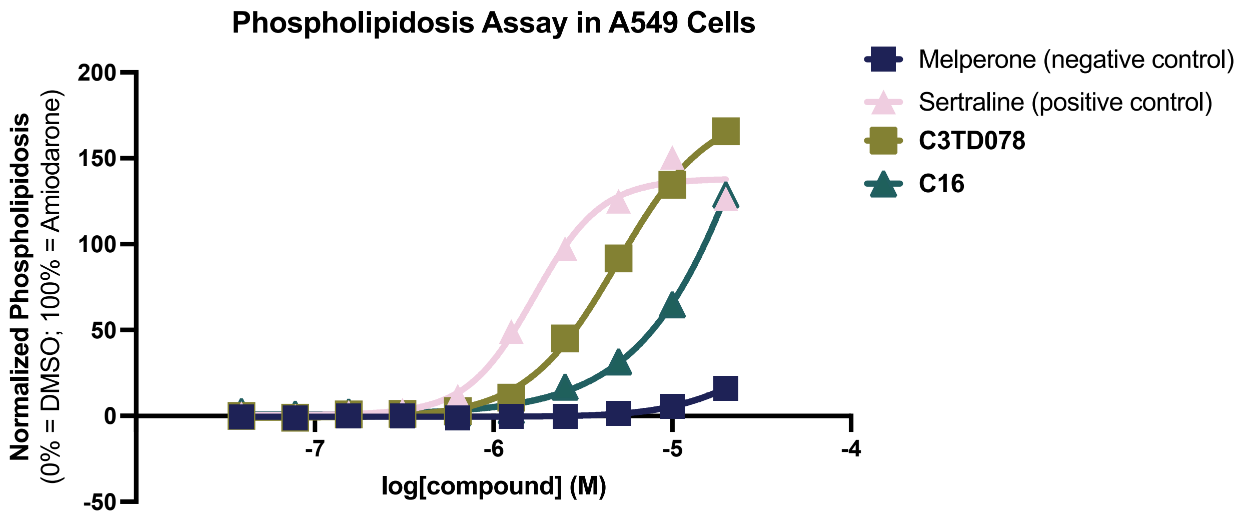


**Supplemental Figure 12.** The ability of **C3TD078** and **C16** to induce phospholipidosis (PLD) was measured using the immunofluorescence assay reported by Tummino *et al.* 2021. Briefly, A549 cells were treated for 24 hours with compounds in the presence of the lipid dye NBD-PE (ThermoFisher #N360), which fluoresces green as it accumulates in vesicular lipid bodies. Total green intensity was measured using a Cytation 5 plate reader and normalized to DMSO-treated (0%) and amiodarone-treated (100%) cells. The positive control sertraline and both **C16**/**C3TD078** induced PLD in A549 cells at doses > 1 μM, with maximum levels at 20 μM compound exceeding 100% of the level induced by amiodarone. A negative control compound, melperone, was unable to induce PLD as expected.

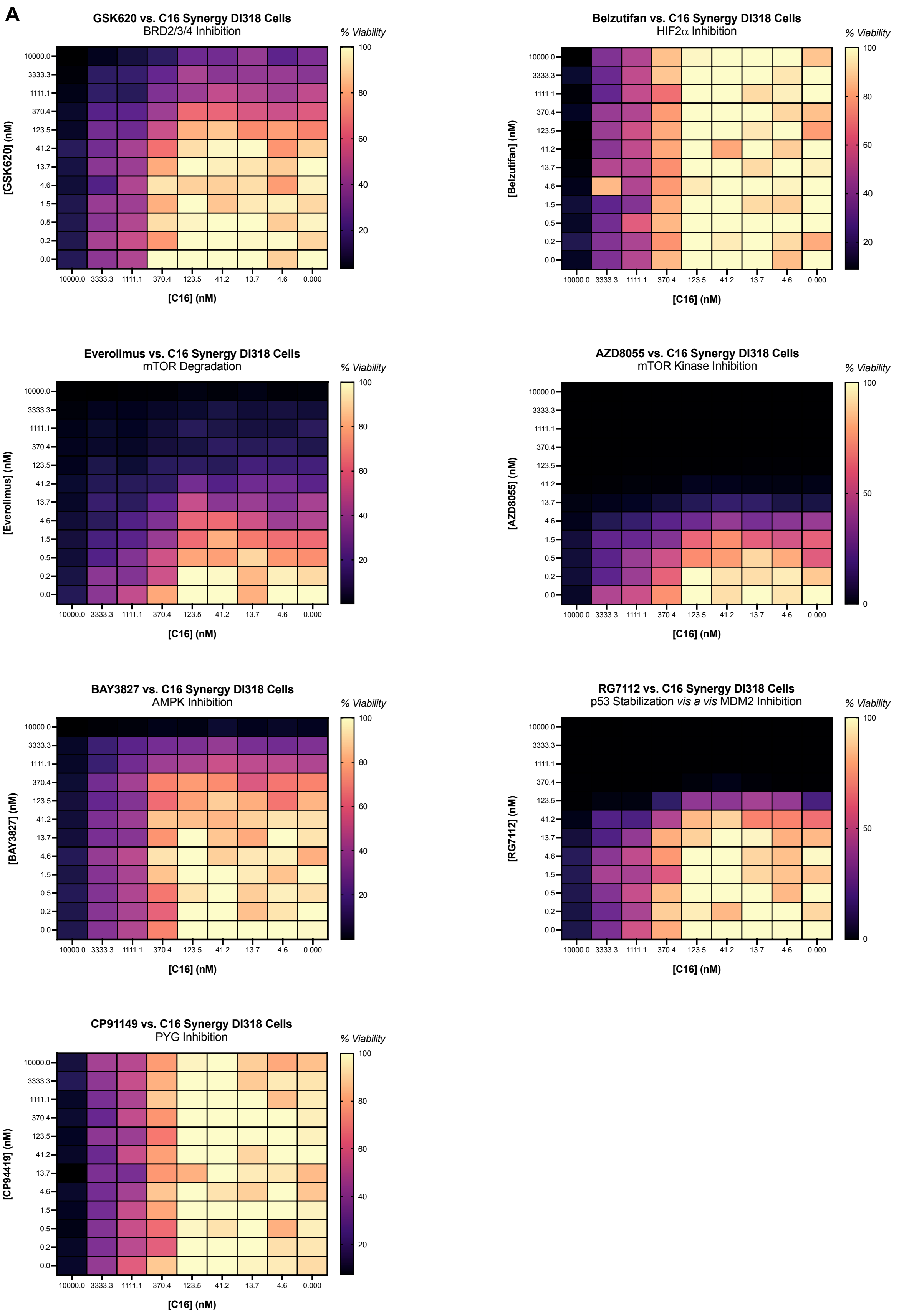
A
GSK620 vs. C16 Synergy DI318 Cells
BRD2/3/4 Inhibition
[GSK620] (nM)
Belzutifan vs. C16 Synergy DI318 Cells
HIF2α Inhibition
[Belzutifan] (nM)
Everolimus vs. C16 Synergy DI318 Cells
mTOR Degradation
[Everolimus] (nM)
AZD8055 vs. C16 Synergy DI318 Cells
mTOR Kinase Inhibition
[AZD8055] (nM)
BAY3827 vs. C16 Synergy DI318 Cells
AMPK Inhibition
[BAY3827] (nM)
RG7112 vs. C16 Synergy DI318 Cells
p53 Stabilization vis a vis MDM2 Inhibition
[RG7112] (nM)
CP91149 vs. C16 Synergy DI318 Cells
PYG Inhibition
[CP94419] (nM)
[C16] (nM)
% Viability
10000.0
3333.3
1111.1
370.4
123.5
41.2
13.7
4.6
1.5
0.5
0.2
0.0
0.000

**B**

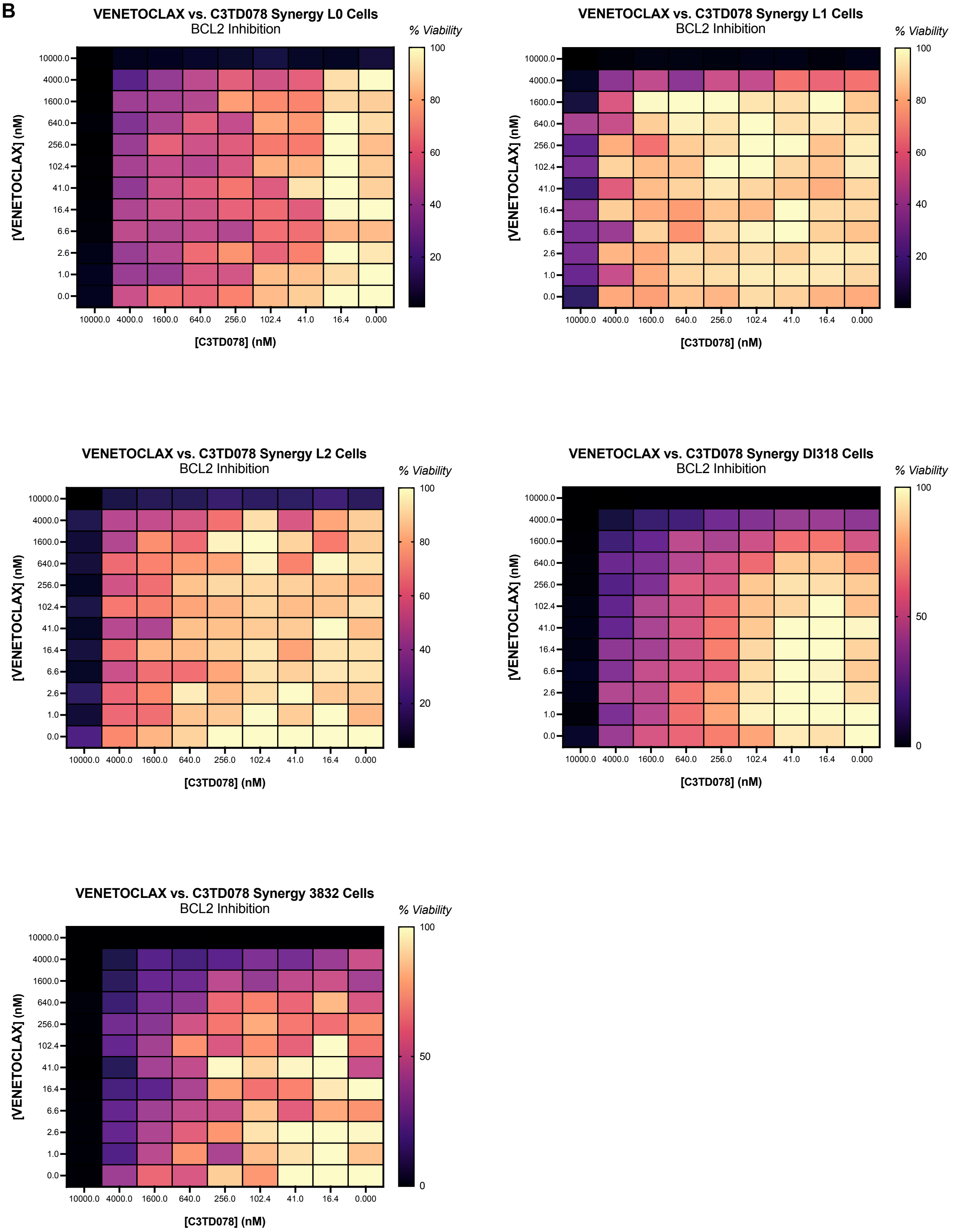


**Supplemental Figure 13. (A)** Cross-titration experiments with **C16** and a variety of rationally selected small-molecules in DI318 CSCs. **(B)** Cross-titration experiments with

**C3TD078** and the BCL2 inhibitor Venetoclax in multiple CSC models. No drug-drug synergy was observed in any combination experiment as estimated by MuSyC (see Materials and Methods). Data are presented as % viability from a single biological replicate after a 7-day treatment period.

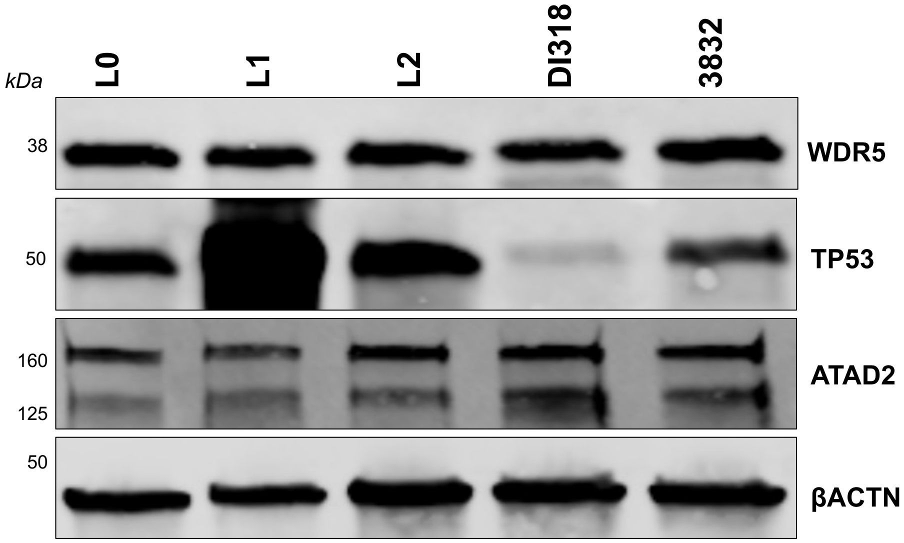


**WDR5:** 1:1000 rb CST1310
**TP53:** 1:500 ms sc-126
**ATAD2:** 1:800 rb CST 50563
**ACTN:** 1:3000 ms CST 3700

**Supplemental Figure 14**. Protein expression of WDR5, ATAD2, and p53 in L0, L1, L2, DI318, and 3832 CSCs as measured by western blot with the indicated antibodies.

A

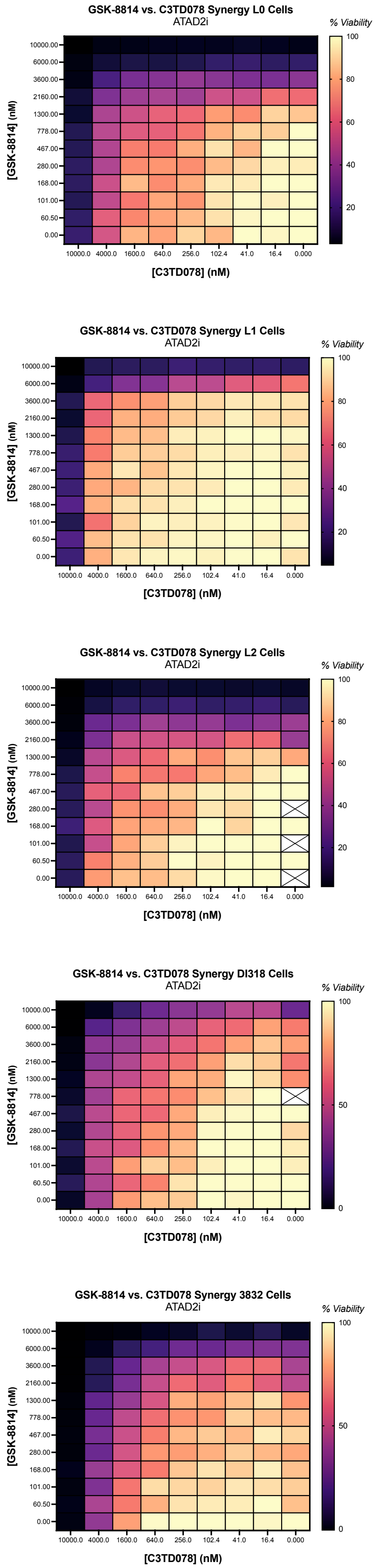
GSK-8814 vs. C3TD078 Synergy L0 Cells
ATAD2i
GSK-8814 vs. C3TD078 Synergy L1 Cells
ATAD2i
GSK-8814 vs. C3TD078 Synergy L2 Cells
ATAD2i
GSK-8814 vs. C3TD078 Synergy DI318 Cells
ATAD2i
GSK-8814 vs. C3TD078 Synergy 3832 Cells
ATAD2i
% Viability
[GSK-8814] (nM)
[C3TD078] (nM)
10000.00
6000.00
3600.00
2160.00
1300.00
778.00
467.00
280.00
168.00
101.00
60.50
0.00
10000.0
4000.0
1600.0
640.0
256.0
102.4
41.0
16.4
0.000

B

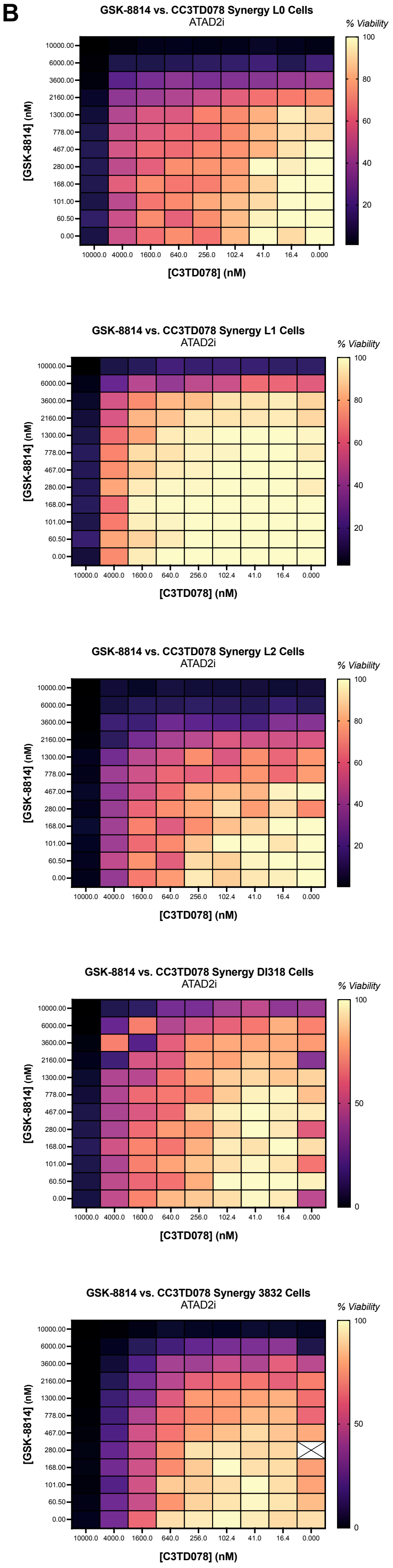
GSK-8814 vs. CC3TD078 Synergy L0 Cells
ATAD2i
GSK-8814 vs. CC3TD078 Synergy L1 Cells
ATAD2i
GSK-8814 vs. CC3TD078 Synergy L2 Cells
ATAD2i
GSK-8814 vs. CC3TD078 Synergy DI318 Cells
ATAD2i
GSK-8814 vs. CC3TD078 Synergy 3832 Cells
ATAD2i
% Viability
[GSK-8814] (nM)
[C3TD078] (nM)
10000.00
6000.00
3600.00
2160.00
1300.00
778.00
467.00
280.00
168.00
101.00
60.50
0.00
10000.0
4000.0
1600.0
640.0
256.0
102.4
41.0
16.4
0.000

**Supplemental Figure 15.** Cross-titration experiments with **C3TD078** and the ATAD2 bromodomain inhibitor GSK-8814 in five different CSC models. Data are presented as % viability from two separate biological replicates, **(A)** and **(B)**, performed as a single technical replicate. No synergy was observed as quantified by MuSyC. Points with an “X” are outliers that have been excluded from the figure for clarity.
